# An estrogen receptor signaling transcriptional program linked to immune evasion in human hormone receptor-positive breast cancer

**DOI:** 10.1101/2024.11.23.619172

**Authors:** Kenichi Shimada, Daniel E. Michaud, Yvonne Xiaoyong Cui, Kelly Zheng, Jonathan Goldberg, Zhenlin Ju, Stuart J. Schnitt, Ricardo Pastorello, Lukas D. Kania, John Hoffer, Jeremy L. Muhlich, Nhan Hyun, Robert Krueger, Alexander Gottlieb, Adam Nelson, Carlos W. Wanderley, Gabriella Antonellis, Sandra S. McAllister, Sara M. Tolaney, Adrienne G. Waks, Rinath Jeselsohn, Peter K. Sorger, Judith Agudo, Elizabeth A. Mittendorf, Jennifer L. Guerriero

## Abstract

T cells are generally sparse in hormone receptor-positive (HR+) breast cancer, potentially due to limited antigen presentation, but the driving mechanisms of low T cell abundance remains unclear. Therefore, we defined and investigated programs (‘gene modules’), related to estrogen receptor signaling (ERS) and immune signaling using bulk and single-cell transcriptome and multiplexed immunofluorescence of breast cancer tissues from multiple clinical sources and human cell lines. The ERS gene module, dominantly expressed in cancer cells, was negatively associated with immune-related gene modules TNFα/NF-κB signaling and type-I interferon (IFN-I) response, which were expressed in distinct stromal and immune cell types, but also, in part, expressed and preserved as a cancer cell-intrinsic mechanisms. Spatial analysis revealed that ERS strongly correlated with reduced T cell infiltration, potentially due to its association with suppression of TNFα/NF-κB-induced angiogenesis and IFN-I-induced HLA expression in macrophages. Preoperative endocrine therapy in ER+/HER2-breast cancer patients produced better responses in ERS-high patients, with TNFα/NF-κB expression associated with reduced ERS. Targeting these pathways may enhance T cell infiltration in HR+ breast cancer patients.

**Statement of Significance:** This study elucidates the immunosuppressive role of ER signaling in breast cancer, highlighting a complex interplay between cancer, stromal, and immune cells and reveals potential approaches to enhance immunogenicity in HR+ breast cancer. These findings offer crucial insights into immune evasion in breast cancer and identify strategies to enhance T cell abundance.

## Introduction

Breast cancer is a heterogeneous disease characterized by variability in tumor size, grade, nodal involvement, and expression of hormone receptors (HR; estrogen receptor (ER) and progesterone receptor) and human epidermal growth factor receptor-2 (HER2), which guide disease stratification and therapeutic decisions (*1–3*). Non-malignant cells within the breast tumor microenvironment (TME) significantly impact the development and progression of the disease (*4, 5*), as well as response to therapy (*6–8*). In particular, tumor infiltrating lymphocytes (TILs), comprised primarily of T cells, play a significant role in shaping therapeutic outcomes. TILs can have both prognostic and predictive value across breast cancer subtypes (*9–11*). Increased TILs are prognostic across triple negative and HER2+ breast cancer (*12*) however are not consistently predictive of responses (*13*), as recent work in HER2+ breast cancer revealed that TILs are prognostic but less so predictive (*14*). However, the role of TILs in HR+ breast cancer is more elusive, potentially due to their much lower frequency compared to other subtypes of breast cancer (*8*).

The extent of T cell infiltration varies widely across breast cancer subtypes and within each subtype there is a range of T cell infiltration (*8*). Triple-negative breast cancer (TNBC) typically exhibits higher levels of T cell infiltration compared to HR+ tumors (*12, 15, 16*) and the abundance of intra-tumoral T cells positively associates with response to both chemotherapy and immune checkpoint blockade (ICB) (*8, 13, 17*). In contrast, HR+ breast cancer is characterized by low T cell numbers and low expression of programmed death-ligand 1 (PD-L1) (*13, 18*), which contribute to its designation as immunologically cold and limited responses to ICB in the metastatic setting (*19, 20*). As anti-PD-1 and PD-L1 therapies work primarily by reinvigorating pre-existing T cells, they are less likely to be effective in tumors with low levels of T cell infiltrate (*21–23*). Indeed, to date, ICB has shown the most success in TNBC patients compared to other subtypes of BC, leading to FDA approvals in patients with PD-L1 positive metastatic TNBC as well as in the neoadjuvant and adjuvant setting for early-stage TNBC (*24–29*). This success has stimulated interest in expanding ICB strategies to HR+ breast cancer, where mechanisms underlying T cell scarcity remain poorly understood.

Recent and emerging data from clinical trials testing ICB in early-stage HR+ breast cancer have provided encouraging results. The I-SPY2 trial showed that adding pembrolizumab to neoadjuvant chemotherapy can improve pathological complete response (pCR) rates in early-stage HR+ breast cancer (*30*). Subsequent trials, including KEYNOTE-756 and Checkmate 7FL demonstrated modest but encouraging improvements in pCR rates with ICB-chemotherapy combinations in high grade primary HR+ breast cancer (*31, 32*). These findings highlight the potential of ICB in HR+ breast cancer, while emphasizing the need for a deeper understanding of the TME and predictors of response in this subtype.

T cell infiltration is influenced by the presentation of peptide antigens on major histocompatibility complex (MHC) molecules, also known as human leukocyte antigen (HLA). Antigen presentation by cancer cells in MHC class-I (MHC-I) complexes is critical for endogenous CD8 T cells to recognize and induce cancer cell killing (*33, 34*). Limited tumor-specific antigens, low HLA expression, or defects in antigen presentation machinery may contribute to low T cell numbers in HR+ breast cancer (*11, 35–38*). Loss or down-regulation of MHC-I expression is a common mechanism of immune evasion in solid tumors (*39–42*) and is particularly relevant in HR+ breast cancer, where lower MHC-I expression was associated with low levels of intratumoral and stromal TILs (*43*). Interestingly, in the Cancer Genome Atlas (TCGA) data (*44*), *HLA-A*, an MHC-I molecule, and *CD8B*, a T-cell specific MHC class coreceptor, were positively correlated with each other but negatively correlated with estrogen receptor 1 (*ESR1*) gene expression (*44, 45*), suggesting a potential role for ER signaling in regulating immune activity. Further insight into the relationship between antigen presentation, T cells and ER signaling may inform effective strategies to induce anti-tumor immune responses in HR+ tumors and/or reveal biomarkers of response to therapy.

This study integrates bulk and single-cell transcriptomics with single-cell multiplexed cyclic immunofluorescence (CyCIF) to investigate the relationship between ER signaling, antigen presentation and T cell activity in HR+ breast cancer. We identified groups of genes – termed ‘modules’ – related to these processes in large-scale datasets (TCGA, METABRIC (*46*)), validated them in independent cohorts, and explored their spatial patterns using CyCIF. We found that the genes related to the ER signaling module were inversely correlated with immune response and T cell activity. Cellular neighborhood (CN) analyses revealed T cell, macrophage, and endothelial cell localization and spatial patterns of MHC/HLA expression linked to ER signaling activity. Using pre-and post-treatment samples from the ACOSOG Z1031B trial (*47, 48*), we showed that endocrine therapy (*47, 48*) suppresses ER signaling and promotes TNFα/NF-κB activity, leading to enhanced immune infiltration in HR+ tumors.

## Results

### Discovery of gene modules linking ER signaling and immunity in hormone receptor-positive/HER2-negative (HR+/HER2-) breast cancer

To determine how ER and immune signaling are related, we first tested the association of key hallmark genes representing these pathways (*ESR1*, *HLA-A*, *CD8A*) in TCGA and METABRIC (*44, 46*) datasets (**Fig. S1A**). *ESR1* was inversely correlated with both *HLA-A* and *CD8A* among HR+/HER2-samples (**Fig. S1B**). This suggests that low T cell numbers in HR+ breast cancer may partly be due to reduced MHC/HLA expression, impaired antigen presentation, and limited immune activation.

Building on this observation, we hypothesized that genes strongly correlated with one another may represent co-regulated biological pathways, and that genes which are associated with *ESR1* expression would be inversely associated with immune signaling. To test this, we curated a list of 778 genes related to ER and immune signaling (**Table S1**) that were analyzed in the TCGA and METABRIC datasets. The RNA-seq-based TCGA dataset provided clearer correlations than the microarray-based METABRIC dataset, so we prioritized TCGA for further analyses and used METABRIC for validation (**Fig. S1C-E**). Pairwise correlation analysis generated three clusters of co-expressing genes: ER signaling (ERS), cell cycle (CC), and immune-related genes (pan-immune) (**Fig. 1A**). The CC cluster showed weak association with ERS, suggesting limited relevance (**Fig. S1F**). Therefore, we focused on the ERS and pan-immune clusters, delineating them into two distinct gene modules: the ERS module (29 genes, including *ESR1*), and the pan-immune module (225 genes, including *HLA-A* and *CD8A*) (**Fig 1A**).

**Figure 1.**
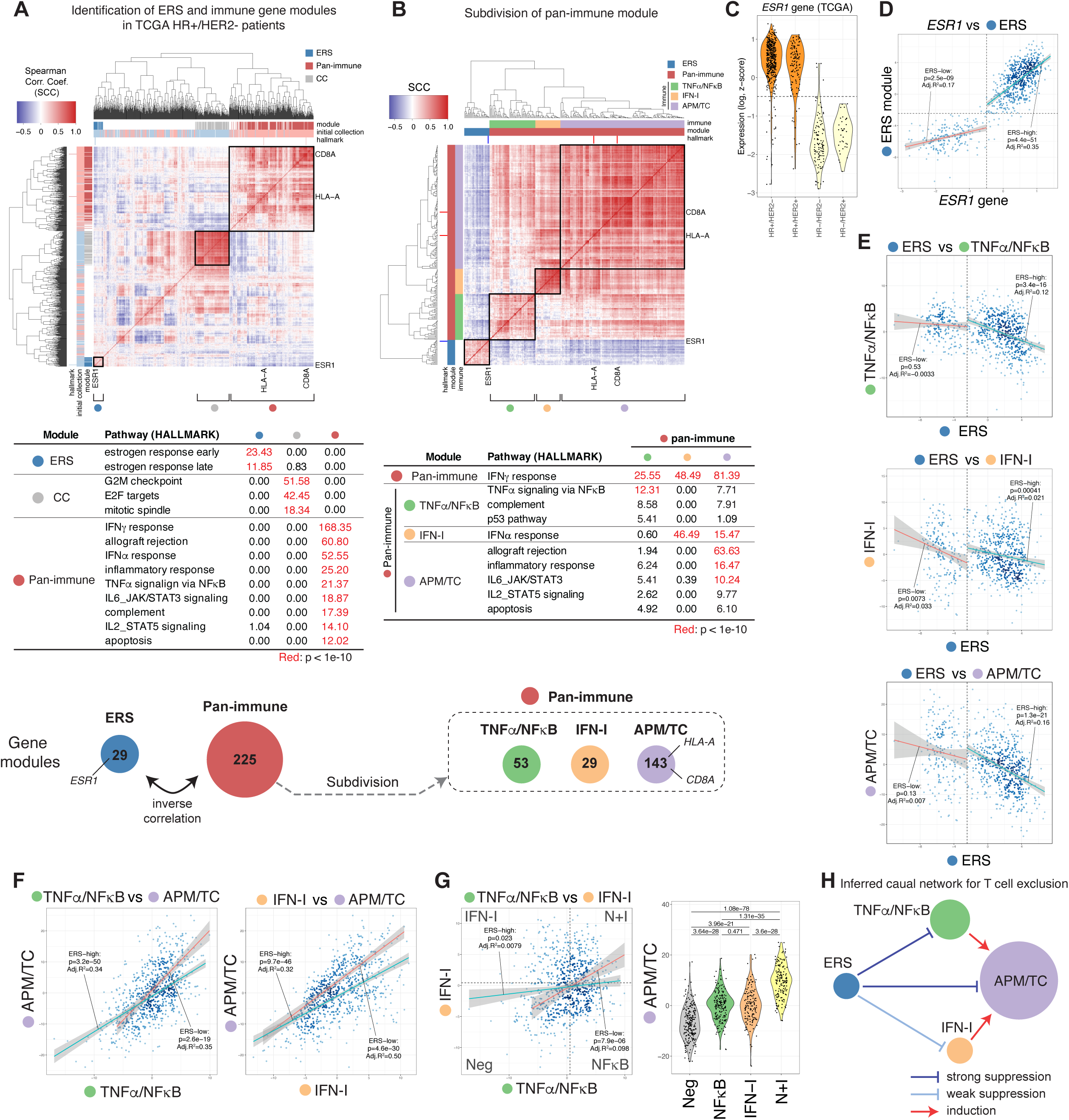
Identification of co-expressed gene modules in HR+ breast cancer patients. **A.** Identification of ERS and pan-immune gene modules from a manually curated gene list. Top: Heatmap displaying Spearman correlation coefficients (SCC) for 778 genes from the manually curated gene list in HR+/HER2-METABRIC patients, with indicators for hallmark genes (*ESR1*, *HLA-A*, *CD8A*), initial gene list (**Table S1**), and final gene modules (ERS, immune, or CC). Squares correspond to correlations within each module. Middle: Enriched hallmark gene sets against each gene module (p < 1e-10, Fisher’s exact test). Bottom: ERS and pan-immune modules that contain 29 and 225 genes respectively, and negatively correlate with each other. **B.** Subdivision of the pan-immune gene module into three distinct modules. Top: SCC for 254 genes that are classified as ERS or pan-immune modules, emphasizing three immune-related modules. The rest of the genes do not belong to robust clusters. Middle: Gene set enrichment across the three immune modules, using the Broad Institute’s hallmark sets with P < 1e-5 from Fisher’s test. Gene sets with P < 1e-10 are highlighted in red. Bottom: The names and the sizes of the three modules. **C.** *ESR1* expression in four breast cancer subtypes from TCGA data. The dashed line corresponds to the *ESR1* expression distinguishing HR+ and HR-samples (AUROC = 0.97). **D.** Analysis of the relationship between *ESR1* gene expression and ERS module expression across patients from all subtypes. Vertical and horizontal lines indicate the thresholds of *ESR1* gene and ERS module, distinguishing their expression in HR+ patients from HR-(AUROC for ERS module is 0.96). Two linear regression lines correspond to *ESR1*-low and *ESR1*-high populations. **E.** Relationship between the expression of ERS module and three immune modules’ expression, differentiated by HR+ versus HR-status. Patients of all subtypes are shown. Two linear regression lines correspond to ERS-low and ERS-high populations. **F.** Relationship of TNFα/NF-κB and IFN-I module expression with APM/TC expression across subtypes. Two linear regression lines correspond to ERS-low and ERS-high populations. **G.** Grouping of patients based on TNFα/NF-κB and IFN-I module expression (left) and corresponding APM/TC expression (right). Two linear regression lines in the left plot correspond to ERS-low and ERS-high populations. The samples were divided into four groups based on their TNFα/NF-κB and IFN-I expression compared with the median values. **H.** Schema showing inferred module interactions illustrating ERS-driven T cell exclusion. Circles represent individual modules and arrows represent inferred causality. Refer to **Figures S1-3** for additional details.

Correlation analysis between the ERS and pan-immune modules reinforced their inverse relationship. While genes within each module were positively correlated, the ERS and immune modules were inversely correlated with each other in HR+/HER2-breast cancer patients (**Fig. 1B**, **Fig. S1G**). The negative correlation aligns with prior work suggesting that ER signaling suppresses T cell responses (*49*). Within the pan-immune module, three distinct clusters emerged, each aligned with unique biological processes derived from the Broad Institute’s ‘hallmark’ gene set (*50*) and were designated as TNFα/NF-κB, IFN-I, and ‘antigen presentation machinery and T cell activity’ (APM/TC) modules (**Table S2**).

### Delineating ERS and immune module interplay across breast cancer subtypes

To determine whether the inverse relationship between ERS and immune modules observed in HR+/HER2-breast cancer is consistent across other breast cancer subtypes, we analyzed the TCGA cohort. Immune-related gene modules (TNFα/NF-κB, IFN-I, and APM/TC) maintained positive correlations with each other across all subtypes, independent of ERS (**Fig. S1G-K**). In contrast, the ERS module exhibited subtype-specific behavior, showing a strong positive correlation within itself and inverse correlation with immune modules only in HR-positive cancers (HR+/HER2-and HR+/HER2+), confirming the HR-dependent nature of this relationship (**Fig. S1G-K**). Across all breast cancer subtypes, ERS module expression was positively correlated with *ESR1* expression (**Fig.1C,D**). Higher ERS expression was linked to reduced TNFα/NF-κB and APM/TC module expression, with this correlation being more pronounced in HR+ samples. The inverse relationship between ERS and immune modules persisted even when HR+ and HR-samples were pooled, with HR+ samples showing higher ERS expression and lower immune module expression compared to HR-samples (**Fig. 1E**).

Among all breast cancer samples, the IFN-I module displayed a weaker correlation with ERS, suggesting nuanced interactions between specific immune pathways and ERS. Further analyses revealed stronger correlations between TNFα/NF-κB and APM/TC, and between IFN-I and APM/TC modules across both HR+ and HR-samples (**Fig. 1F**). Notably, TNFα/NF-κB and IFN-I modules showed minimal correlation with each other (**Fig. 1G**), suggesting distinct biological processes govern their activity. These observations may have important implications for understanding how ER signaling influences immune suppression through specific immune pathways (*51*).

Our findings were validated using the METABRIC dataset and an independent cohort of 74 primary breast cancer patients from the University of Texas MD Anderson Cancer Center. Consistent correlations of the gene modules across datasets and subtypes support their robustness (**Fig. S2, S3**). Together, these analyses suggest that ER signaling exerts differential suppression of immune responses, with the strongest inverse correlation observed with TNFα/NF-κB and APM/TC modules, and a weaker association with IFN-I (**Fig 1H**). The differential activity of TNFα/NF-κB and IFN-I modules, despite their shared correlation with APM/TC, underscores their distinct biological roles (*52–58*).

### Unraveling gene module expression at the patient and individual cell level using single-cell transcriptomic analysis

To explore whether differences in module correlations observed in bulk sequencing reflect cell type-specific expression patterns, we analyzed a single-cell RNA sequencing (scRNA-seq) dataset of 34 treatment-naïve primary breast cancers spanning various subtypes (20 HR+/HER2-, 6 HER2+, 8 TNBC) from the Walter and Eliza Hall Institute of Medical Research (WEHI) (*59*). Cell types were identified using lineage-specific markers (**Fig. 2A**, **Fig. S4**). Consistent with bulk analyses, clustering analysis revealed co-regulation among genes within the same modules (**Fig. S5A,B**).

**Figure 2.**
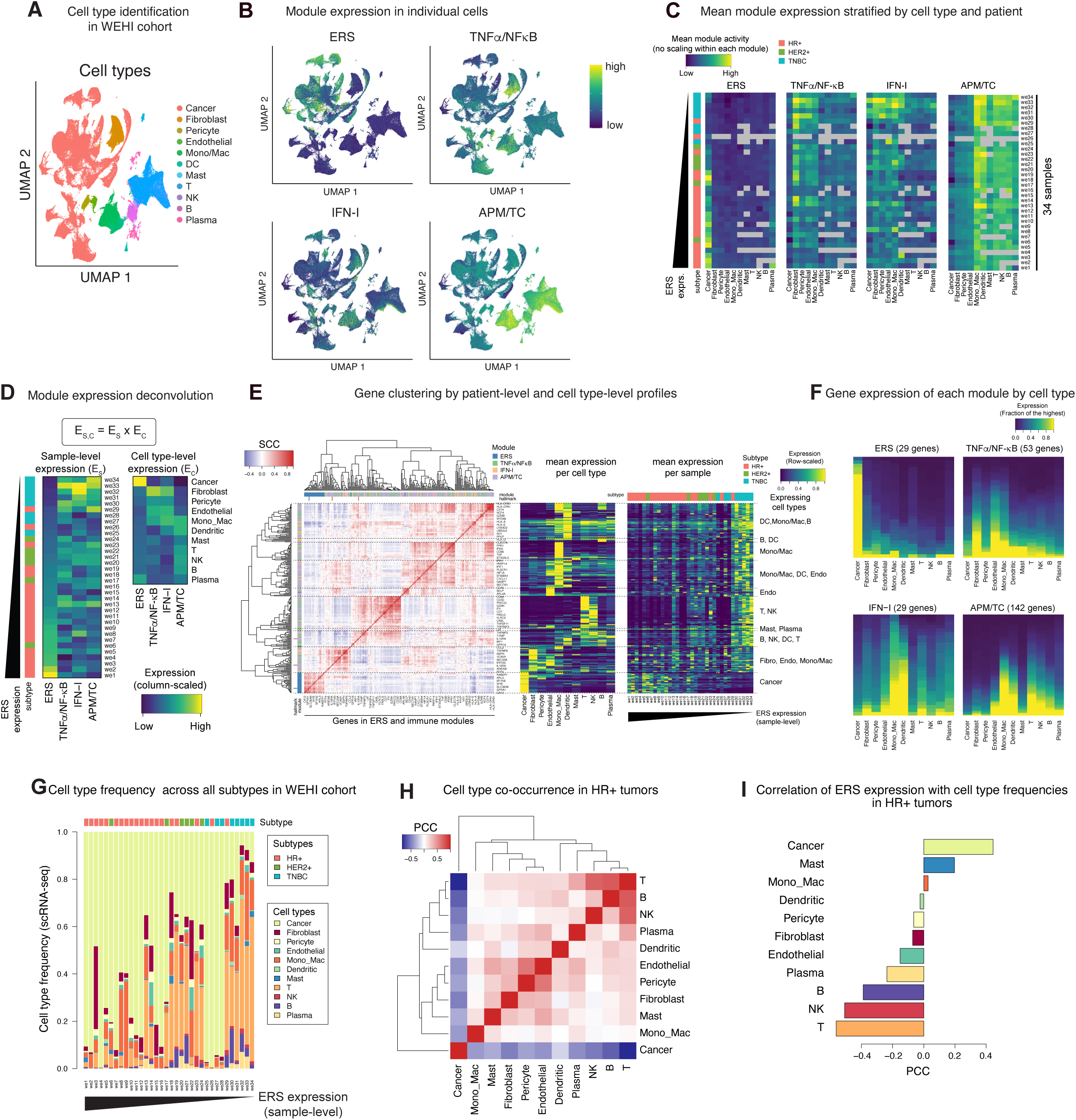
scRNA-seq data of primary breast cancer reveals patient level and cell type level module expression. **A.** Identification of cell types in scRNA-seq data from 34 treatment-naïve HR+ breast cancer patients in WEHI cohort, using lineage marker expression. **B.** UMAP representation of individual cells showing gene module expression, assessed using ssGSEA enrichment scores. **C.** Heatmap of each module’s mean expression (ssGSEA enrichment scores) by cell types and by samples. Each module’s mean expression was plotted by cell type (x-axis) and patient (y-axis) and the resulting heatmaps demonstrate variation in module gene expression by both patient and cell type. **D.** Module expression was represented at the patient level (left) and cell type level (right, transposed), which were deconvolved from mean ssGSEA enrichment score matrices in **C** using the mathematical model on the top. Samples in **C** and **D** are in the same order, sorted by the patient-level ERS expression. **E.** Clustering of genes in the four modules based on their per-cell type and per-patient level profile. Pearson correlation analysis of the per-cell type and per-patient level profile (left); the expression per cell type is averaged over all patients (middle), and the expression per patient is averaged over cell types (right). **F.** Heatmap of gene expression by cell type for each module. The middle panel from **2E** is reorganized and displayed separately for each module. The gene expression is shown a the fraction of the highest expressing cell type (0-1) and are sorted within each column to emphasize the cell type’s contribution. Note that rows are not comparable across cell types within each module. **G.** Overview of cell type frequency of all 34 samples. **H.** Cell type co-occurrence assessed by PCC across 20 HR+ samples. **I.** Correlation of ERS module expression per patient with each cell type’s frequency. The analysis is done among 20 HR+ samples. Refer to **Figures S4-5** for additional details.

Module expression in individual cells was computed using single-sample GSEA (ssGSEA) enrichment scores (*60*). UMAP visualization showed that each gene module was predominantly expressed in distinct cell populations. ERS module expression was highest in luminal cancer cells with low custom ‘basality’ scores (**Fig. 2B**, **Fig. S4A**), and *ESR1* gene expression was positively correlated with ERS module expression in cancer cells, confirming the relationship observed in bulk analyses (**Fig. S5A**). Immune-related modules (TNFα/NF-κB, IFN-I, and APM/TC) showed distinct expression patterns across cell types and varied among patients (**Fig. 2C**). Module expression analysis across cell types and patients revealed a strong inverse correlation between ERS and immune modules across patients, aligning with findings from bulk data (**Fig. 2C**). Further analysis of modules across cell types revealed that each module’s expression was more consistent across cell types than between different modules (**Fig. S5B**). This prompted the use of matrix decomposition to separate module expression into patient-level and cell type-level components (**Fig. 2D**). ERS module expression was consistently localized to cancer cells, while immune modules were expressed primarily in non-cancer cells. Specifically, TNFα/NF-κB module genes were highly expressed in fibroblasts and endothelial cells, IFN-I module genes in monocytes and macrophages, and APM/TC module genes were primarily expressed in lymphocytes and myeloid cells (**Fig. 2D-F**).

The inverse correlation between ERS and immune modules was confirmed at the patient level across all breast cancer subtypes in the WEHI cohort and was validated using the subset of 20 HR+ patients in the same cohort (**Fig. S5C**). Taken together, these data demonstrate that gene module expression reflects both patient-level variability and cell type-level activities. To investigate how the expression of each module is influenced by both cell type-specific expression patterns and differences in cell type frequencies may contribute to variability in module expression observed in bulk analyses, we analyzed cell type frequencies across the WEHI cohort. Stromal and immune cell frequencies varied widely, illustrating a diverse and complex TME (**Fig. 2G**). Correlation analysis revealed that cancer cell abundance negatively correlated with immune cell abundance, especially lymphoid cells (T, NK, B, and plasma cells) and dendritic cells. (**Fig. 2H**, **Fig. S5D**). This trend, although influenced partly by normalization to total cell counts, suggests that increased cancer cell prevalence is associated with diminished stromal and immune cells in the TME. Notably, ERS gene module expression was positively correlated with cancer cell abundance and negatively with T cell infiltration, linking ERS activity to reduced T cell presence in HR+ breast cancer (**Fig. 2I**, **Fig. S5E**).

### Cancer cell-intrinsic relationship between ERS and immune modules in breast cancer cell lines

Our prior analyses presented here established that ERS is predominantly expressed in cancer cells, while immune gene modules are primarily expressed in non-cancer cells. However, cancer cells also exhibited expression of immune modules, albeit, at lower levels (**Fig. 2C,D**), suggesting a cancer cell-intrinsic component to immune signaling that may impact TME interactions. To test whether the inverse relationship between ERS and immune modules observed in primary tumors is preserved within cancer cells, we analyzed RNA-seq data from 54 breast cancer cell lines in the Cancer Cell Line Encyclopedia (CCLE; **Fig. 3A**), spanning various molecular subtypes classified by PAM50 (*61, 62*). We aimed to distinguish cancer cell-intrinsic modules, preserved within cancer cells, from the whole-tumor modules defined in bulk transcriptome analyses. We revealed that gene expression patterns in CCLE cell lines closely mirrored those in cancer cells from the WEHI cohort, with 61% of the total number of identified module genes (153 genes) expressed in the cell lines (**Fig. S6A-B**). Notably, 76-97% of genes in the TNFα/NF-κB and IFN-I modules were detected in cell lines, compared to only 41% of APM/TC genes, which are predominantly associated with lymphoid cells (**Fig. 3B**). Correlation analysis within CCLE data, conducted both within the 19 luminal-specific cell lines (classified using PAM50 and characterized by HR expression and a more differentiated phenotype) and also across all 54 cell lines, revealed three distinct clusters of co-expressed genes – ERS, TNFα/NF-κB, and IFN-I – strongly overlap with the gene modules identified in bulk analyses (**Fig. 3C-E**, **Fig. S6C-D**). APM/TC genes did not form a distinct cluster on its own; however, some APM/TC genes clustered with TNFα/NF-κB and IFN-I genes. These findings suggest that the TNFα/NF-κB and IFN-I modules derived from whole tumors contain subsets of genes with preserved expression in cancer cells, referred to as cancer cell-intrinsic expression.

**Figure 3.**
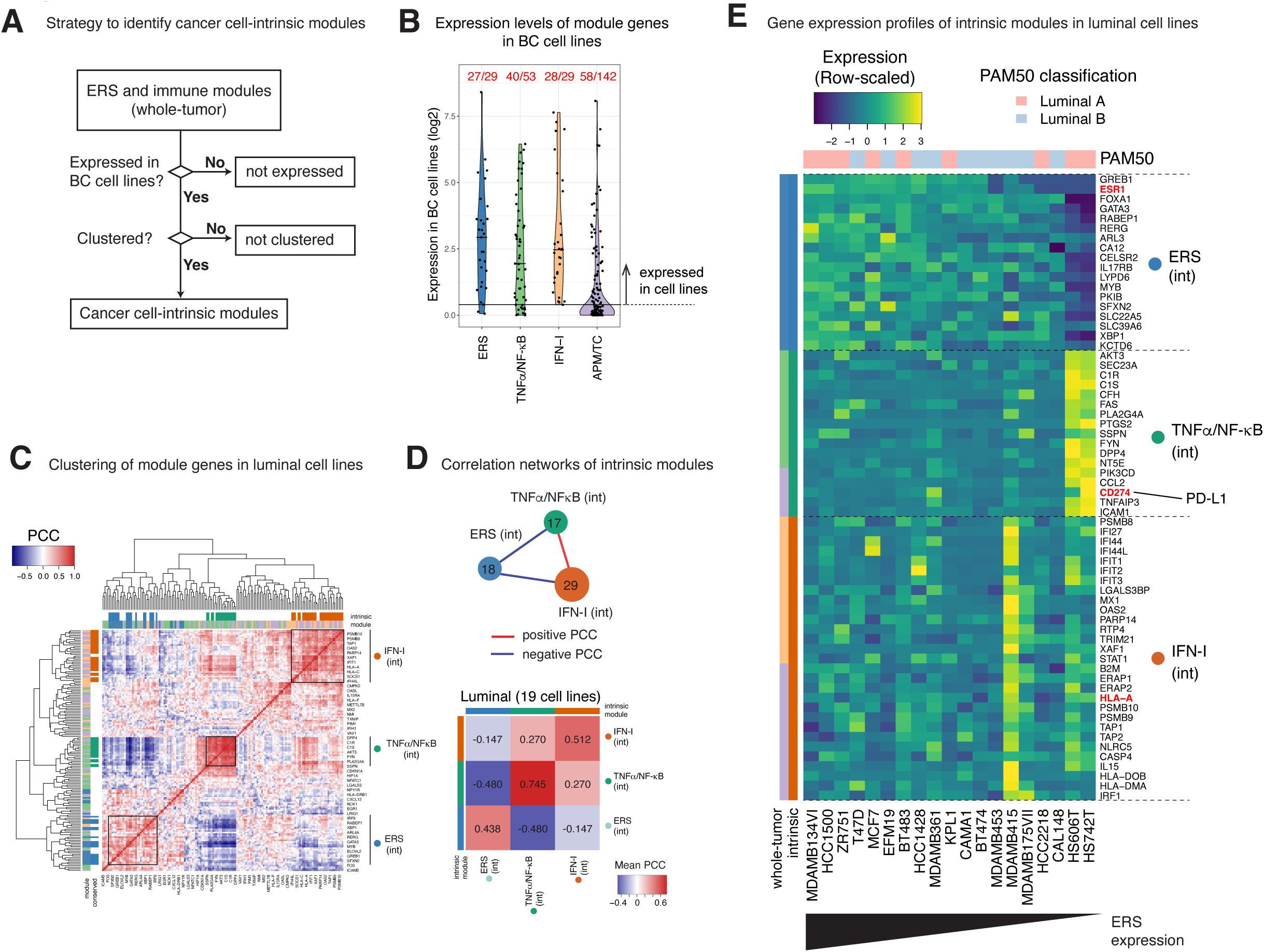
Co-regulation of ERS and immune modules are preserved within breast cancer cell lines. **A.** The process to identify the gene module subsets whose co-expressing relationship is preserved in breast cancer cell lines. **B.** Violin plot showing expression levels of individual genes in each module are expressed in 54 breast cancer cell lines. Each point represents the expression of a single gene, with the horizontal line indicating the signal threshold computed in **Fig. S6B**; genes below this line are considered not expressed in cell lines. The fractions at the top indicate the number of genes expressed in cell lines divided by the total number of genes in each module. Most genes expressed in T or NK cells in the WEHI cohort were not expressed in cancer cells from the cohort or in cell lines. These genes, primarily from the APM/TC module, are below the threshold. **C.** Hierarchical clustering of module genes expressed in breast cancer cell lines, based on their expression patterns across 19 PAM50 luminal breast cancer cell lines. Three cancer cell-intrinsic modules (ERS, TNFα/NF-κB, IFN-I) are highlighted with squares and annotated on the right. **D.** Network diagram illustrating the interactions between three intrinsic modules in breast cancer cell lines (top); Heatmap depicting the mean PCC for each module derived from the data in **C** (down). **E.** Individual gene expression patterns of cancer cell-intrinsic ERS, TNFα/NF-κB, and IFN-I modules in 19 luminal cell lines. The intrinsic IFN-I gene module was further subdivided into, IFN_sub and APM_sub, with pathway enrichment highlighted (see **Fig. S6F,G** for pathway analysis). Refer to **Figure S6** for additional details.

The cancer cell-intrinsic TNFα/NF-κB module was enriched for genes in the complement pathway and included PD-L1 (*CD274*), suggesting that immune activity within this module may drive PD-L1 upregulation. Its upregulation could contribute to T cell exhaustion through the PD-L1–PD-1 axis (**Fig. 3E**). Similarly, the intrinsic IFN-I module was linked to both type I interferon responses and antigen processing and presentation, with distinct contributions from IFN-I (termed ‘IFN_sub’) and APM/TC (‘APM_sub’) genes (**Fig. S6E-G)**. Both TNFα/NF-κB and IFN-I intrinsic modules were overexpressed in basal cell lines, which are linked to more aggressive and undifferentiated phenotypes. The basal cell lines showed minimal expression of ERS module genes, including *ESR1* (**Fig. S6H**).

### Module expression and cell type frequency in HR+ breast cancer

To investigate the relationship between ERS and immune gene module expression within specific cell types related to their spatial distribution, we integrated bulk RNA-seq, single-nucleus RNA sequencing (snRNA-seq), and multiplexed CyCIF (*63–67*) on 29 primary HR+ breast cancer samples from the Dana-Farber/Brigham Cancer Center (DF/BCC) (**Table S3**).

The bulk transcriptomic analysis of the DF/BCC cohort corroborated the inverse correlation between ERS and immune gene module expression observed in the TCGA and METABRIC analyses (**Fig. S7A-C**). snRNA-seq data further confirmed the distinct expression patterns, with the ERS module genes predominantly expressed in cancer cells, while immune-related modules were expressed in stromal and immune cells (**Fig. S7D-J**).Consistent with prior findings, cell type frequency analysis revealed that increased cancer cell prevalence was associated with decreased immune and stromal cell prevalence, confirming an inverse correlation between cancer cells and non-cancer cells in the TME (**Fig. S7K-M**). While normalization to cell proportions within each sample partly influences this relationship, the data demonstrate a consistent trend where higher cancer cell abundance corresponds to lower immune cell frequency.

To directly visualize cellular composition, CyCIF was used to identify individual cells in tumor sections corresponding to the same tumors analyzed by bulk RNA-seq and snRNA-seq (**Fig. S8**). CyCIF analysis confirmed a strong positive association between cancer cell abundance and ERS module expression, and an inverse association with immune gene modules (**Fig. 4A,B**). CyCIF analysis identified five specific cell types: CD8+ T cells (CD8T; CD3+/CD8+), regulatory T cells (CD4T_Treg; CD3+/CD4+/FoxP3+), non-Treg CD4+ T cells (CD4T_nonTreg; CD3+/CD4+/FoxP3-), Mac_CD163 (CD68-/CD163+), and Mac_CD68_CD163 (CD68+/CD163+) that consistently co-occurred in ERS-low samples (**Fig. 4B,C**). These cell types were often found together and were enriched in samples with higher infiltration of CD4+ and CD8+ T cells, both locally and globally, in subsequent analyses (e.g., **Fig. 5A,B**, **6E**). Notably, TNFα/NF-κB and IFN-I gene module activities derived from bulk transcriptomic analysis correlated most significantly with the abundance of endothelial cells and CD163+ macrophages identified by CyCIF, suggesting specific roles for these cell types in the TME of HR+ breast cancer (**Fig. 4C**).

**Figure 4.**
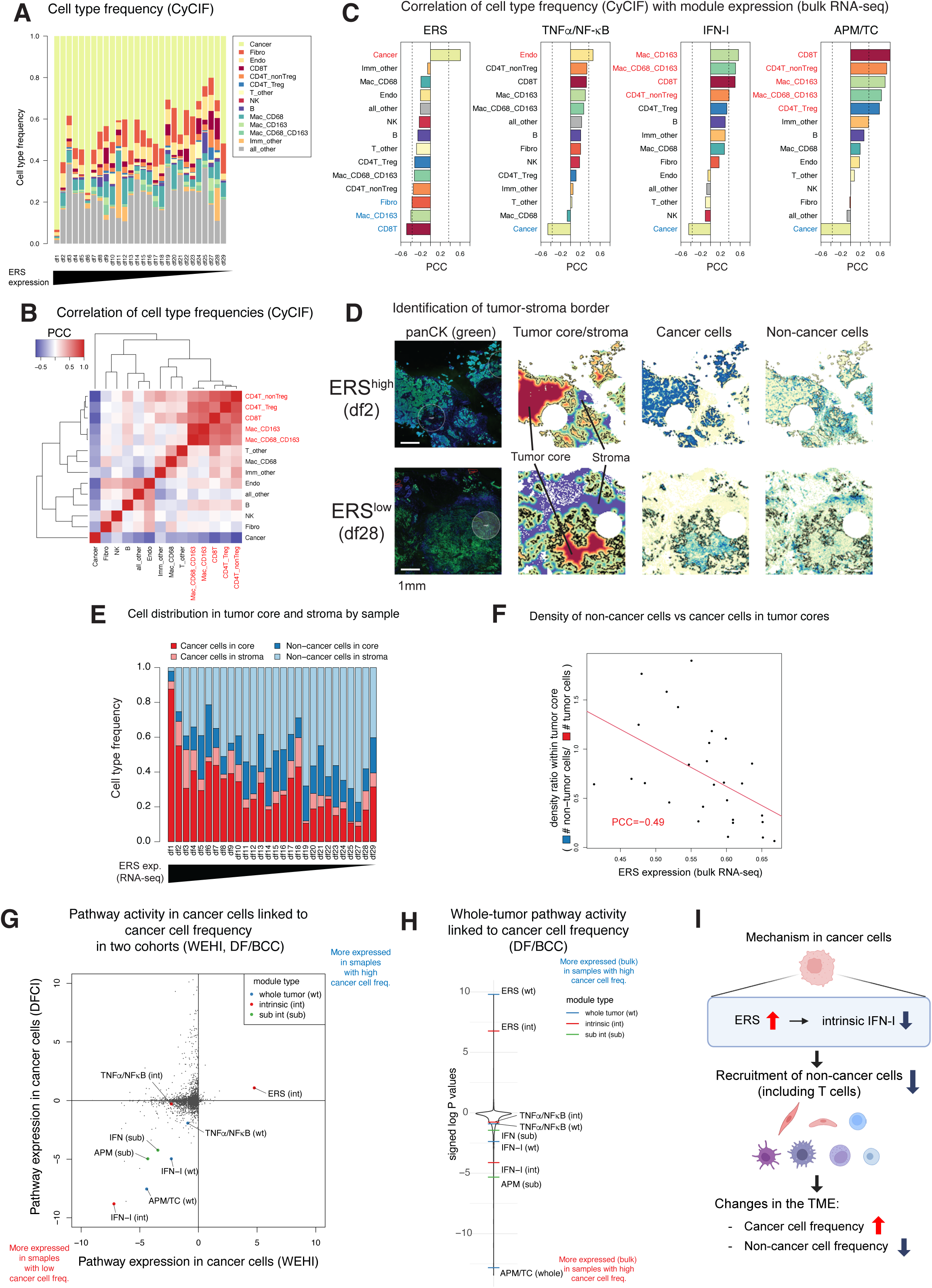
Transcriptional module expression is correlated with corresponding cell type frequency in HR+ breast cancer. **A.** Cell type frequency of 28 samples reconstituted from the CyCIF data. **B.** Cell type co-occurrence across 28 patients analyzed from the CyCIF data. **C.** Pearson correlation coefficients between the four modules’ expression and cell type frequency reconstituted from the CyCIF data. Cell types highlighted in red and blue are significantly positively and negatively correlated, respectively, with the corresponding module expression (p-value < 0.05 in Pearson correlation test). **D.** Computational identification of the tumor-stromal border in CyCIF data analysis. Two examples (ERS expression-high and low tumors) are shown, featuring images with panCK staining, computationally identified tumor core and stromal regions, and the density of tumor and non-cancer cells from 20µm-radius cellular neighborhoods, in the same region of interest (ROI). **E.** The frequency of tumor and non-cancer cells within the tumor core and stroma for each sample. **F.** Relationship between the ratio of the density of non-cancer cells to cancer cells within the tumor core compared to their ERS module expression. **G.** Pathway activity in cancer cells plotted against their association with cancer cell frequency in the WEHI cohort (x-axis) and the DF/BCC cohort (y-axis). Whole-tumor modules (including both intrinsic and extrinsic) are highlighted in blue, intrinsic in red, and intrinsic module subsets in green. **H.** Pathway activity in the whole tumor, assessed by bulk RNA-seq, and its association with cancer cell frequency. **I.** Schematic diagram describing the suggested mechanism of non-cancer cell exclusion induced by ER signaling. Refer to **Figure S7-8** for additional details.

**Figure 5.**
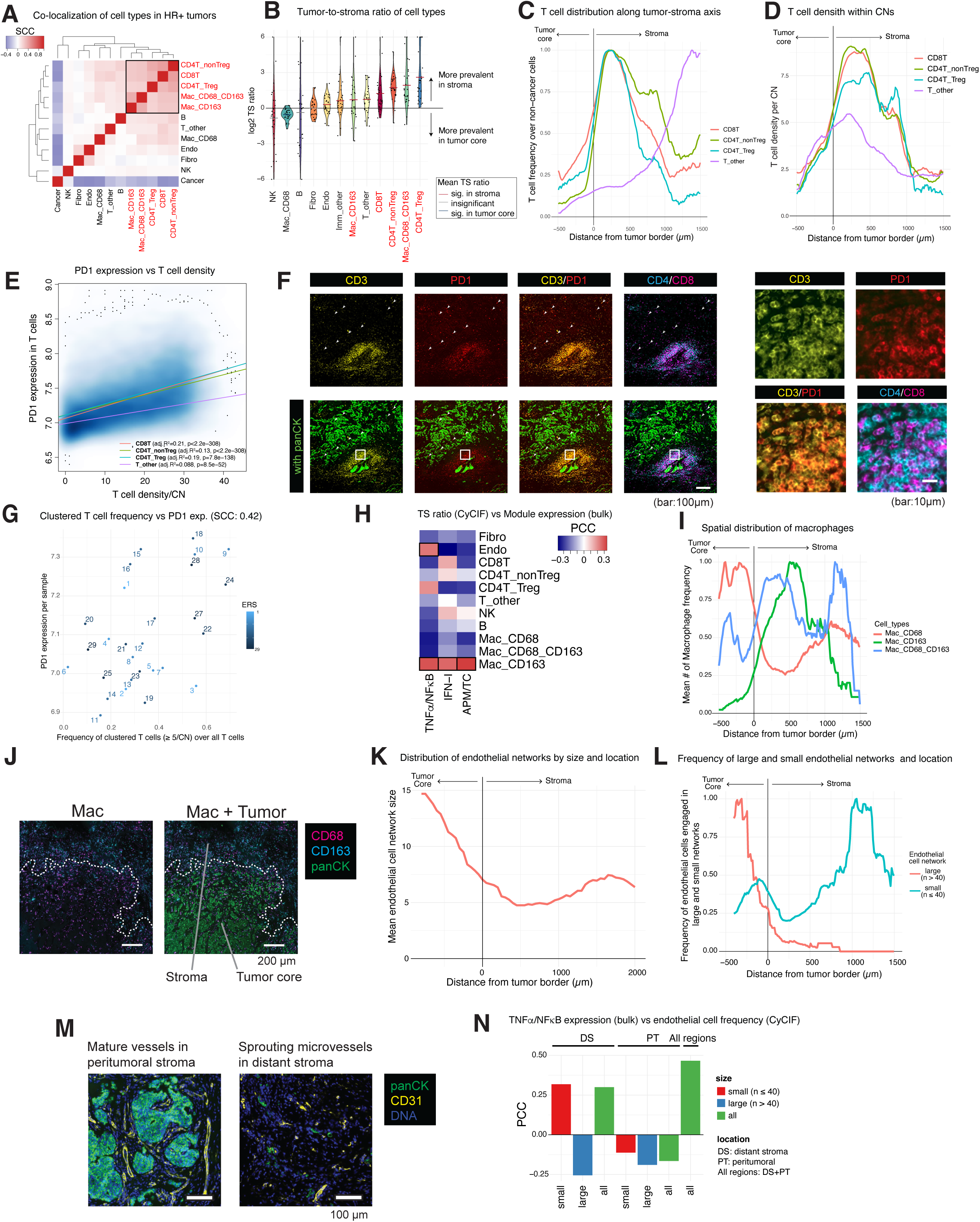
Spatial analysis of T Cells in HR+ breast cancer highlights the impact of ERS module expression. **A.** Co-localization analysis of cell types among 28 samples, assessed by Spearman correlation of cell type frequency across CNs. **B.** Tumor-to-stroma ratio of each cell type across 28 samples. **C.** Spatial distribution of T cells along the tumor-stroma axis. T cell frequency, normalized relative to non-cancer cells at each distance, is scaled so that the maximum value is set to one. **D.** Average T cell density in a 20µm cellular neighborhood along the tumor-stroma axis. **E.** Relationship between PD-1 expression and the local T cell density per CN for each T cell subtype. **F.** Representative images of PD-1+ clustered and non-clustered T cells. Arrowheads show PD-1+ non-clustered T cells within tumor core. Magnified views of the area highlighted by the square in the bottom row are shown on the right. **G.** Scatterplot between the proportion of T cells engaged in clustered T cells and mean PD-1 expression across 28 samples. **H.** Relationship between the tumor-stromal ratio of each cell type and each module expression across 28 samples. A positive PCC indicates the greater stromal prevalence of a cell type in samples with a higher module expression. **I.** Spatial distribution of three subtypes of macrophages along the tumor-stroma axis. Macrophage frequency, normalized relative to non-cancer cells at each distance, is scaled so that the maximum value is set to one. **J.** Representative images highlighting distinct spatial distributions of Mac_CD68 and Mac_CD163 concerning the tumor-stromal border. **K.** Spatial distribution of endothelial network size (number of physically connected endothelial cells), with mean network size is calculated along the distance from tumor border. **L.** Spatial distribution of endothelial cells along the tumor-stroma axis, stratified by the size of the physically connected networks—large (n > 40) or small (n ≤ 40). Their frequency among non-cancer cells is normalized so the peak is set to one. **M.** Representative images highlighting large and small vascular networks. **N.** Association between endothelial cell network size and location (distant stroma, peritumoral, or combined) and TNFα/NF-κB expression across 28 samples. In this analysis, distant stroma (DS) refers to regions beyond 500 µm from the tumor border (d > 500), while peritumoral (PT) refers to areas within 500 µm, including tumor core. All regions include both DS and PT. The frequency of endothelial cells compared to non-cancer cells in each location is evaluated. A high positive PCC indicates a strong correlation between endothelial cell abundance and higher TNFα/NF-κB expression. Refer to **Figure S9** for additional details.

### Spatial exclusion of non-cancer cells in tumors with high ERS module expression

To investigate whether ERS and immune gene module expression are associated with spatial organization in HR+ breast cancers, we classified individual cells based on their proximity to the tumor core, defined in CyCIF images as clusters of ≥10 physically connected cancer cells. Areas outside the tumor core were designated as stroma, while the tumor border was defined as the peripheral edge of these connected clusters (**Fig. 4D**). Our analysis revealed that ERS-high samples had a significantly lower frequency of non-cancer cells (e.g., stromal and immune cells), particularly in the tumor core, compared to ERS-low samples (**Fig. 4E**). Non-cancer cell density was markedly lower in the tumor core of ERS-high samples, suggesting a spatial exclusion of non-cancer cells in regions of high ERS activity (**Fig. 4F**).

Across two independent cohorts (WEHI and DF/BCC), we consistently observed that high ERS module expression in cancer cells correlated with increased cancer cell frequency and decreased infiltration of non-cancer cells. The reduced infiltration of non-cancer cells in ERS-high samples suggests that ERS activity in cancer cells may suppress the recruitment of non-cancer cells to the tumor core. We further identified that high cancer cell frequency was strongly linked to suppression of the cancer cell-intrinsic IFN-I module (**Fig. 4G**). Importantly, this intrinsic IFN-I module emerged as the pathway most strongly associated with cancer cell frequency among over 5,000 analyzed pathways (**Fig. 4H**). However, at the whole-tumor level, bulk RNA-seq data revealed that suppression of the APM/TC module correlated more strongly with cancer cell frequency compared to the intrinsic IFN-I module, likely due to its broader role in mediating infiltrating immune cells (**Fig. 4I**). Taken together, these data suggest that suppression of the intrinsic IFN-I module within cancer cells mediates the connection between ERS activity and immune exclusion in HR+ breast cancer (**Fig. 4J**).

### Spatial localization of T cell and macrophage subsets in the stroma

To investigate the impact of ER signaling on tumor architecture, we analyzed individual stromal and immune cells in greater detail, which, in part, is available as an interactive web tool using Minerva (*68*) in **Supplementary Data S1-2**. Initial co-localization analysis within cellular neighborhoods (CNs) revealed that CD4+ and CD8+ T cell subsets frequently co-localized within 20µm-radius CNs, often in proximity to macrophage populations Mac_CD68_CD163 and Mac_CD163 (**Fig. 5A**, **Fig. S9A**). These T cell and macrophage subsets were more abundant in samples with higher APM/TC module expression at the bulk level and were predominantly located in the stroma compared to the tumor core (**Fig. 5B**). CN analysis further confirmed that CD4+ (both Treg and non-Treg) and CD8+ T cells clustered near the tumor border, with their densities peaking within 500 µm of the tumor edge (**Fig. 5C**). By contrast, T_other cells displayed distinct spatial patterns, appearing more frequently in the distant stroma rather than clustering near the tumor core (**Fig. 5C,D**, **Data S1-2**). Other lymphocyte populations, such as NK and B cells, showed inconsistent spatial distribution across samples and no significant association with ERS. As a result, their link to ERS activity remains unclear. Therefore, we prioritized CD4+ and CD8+ T cell subsets, along with macrophage populations, for further investigation.

### PD-1 protein expression on T cells is associated with the local T cell density

To further explore the spatial distribution and functional state of T cells, we assessed PD-1 protein expression. PD-1 expression was highest in T cells located within the tumor core (**Fig. S9B**), though these cells were relatively rare(**Fig. 5C**). Importantly, PD-1 expression was more strongly linked to the local density of T cells within CNs than to their physical distance from the tumor border, measured as signed distance along the tumor-stroma axis. This suggests that clustered T cells are more likely to be PD-1+ compared to non-clustered ones (**Fig. 5E**). This pattern was consistent across all T cell subsets, especially CD4+ and CD8+ T cells, with clusters predominantly located in the peritumoral stroma, within 500 µm of the tumor border (**Fig. 5D**, **Data S1-2**). Notably, T cell density (count/mm²) showed a weaker correlation with mean PD-1 expression (SCC=0.30), whereas the proportion of aggregated T cells among total T cells showed a stronger correlation (SCC=0.42; **Fig. 5F,G**, **Fig. S9C**). These findings confirm that PD-1 expression is enriched in clustered T cells near the tumor border.

### Roles of stromal Mac_CD163 and endothelial cells in ERS-low tumors

Macrophages expressing CD163 (Mac_CD163) were significantly associated with increased expression of all three immune modules (**Fig. 5H**, **Fig. S9D**). Mac_CD163 and Mac_CD68_CD163 macrophages were more prevalent in the stroma, whereas Mac_CD68 macrophages were predominantly localized in the tumor core (**Fig. 5B,I,J**, **Fig. S9E, Data S1-2**). Taken together with association between cell type frequencies and module expression (**Fig. 4C**) and spatial distribution (**Fig. 5I**), these findings suggest a strong link between stromal Mac_CD163 and Mac_CD68_CD163 macrophages and IFN-I and APM/TC expression in HR+ breast cancers (**Fig. 5A,B,H**). The presence of Mac_CD163 and Mac_CD68_CD163 in the stroma, along with their colocalization with PD-1+ T cells, may indicate a mechanism of suppression in HR+ breast tumors, especially given their abundance in the stroma of ERS-low samples.

Endothelial cells were identified to be more abundant in the stroma of samples with increased TNFα/NF-κB module expression (**Fig. 5B,H**, **Fig. S9F**). These cells formed larger vascular networks, consisting of ≥ 40 physically connected endothelial cells within the tumor core and peri-tumoral stroma, with network size diminishing further from the tumor border (**Fig. 5K-M**). Smaller vascular networks, defined by < 40 endothelial cells, were more common in the stroma distant from the tumor border (> 500 µm), and appeared to represent immature endothelial sprouts rather than functional blood vessels. These smaller networks likely represent early-stage pathological angiogenesis (*69–71*).

The abundance of small vascular networks in the distant stroma, beyond 500 µm from the tumor border, showed a positive correlation with TNFα/NF-κB module expression, while small networks within the stroma or large networks in both stroma and tumor core exhibited a weakly negative association with TNFα/NF-κB module expression (**Fig. 5N**, **Data S1-2**). This suggests that TNFα/NF-κB may regulate stromal microvascular architecture, influencing immune cell trafficking. In ERS-low samples, the association between TNFα/NF-κB activity and stromal endothelial networks highlights its potential role in modulating conditions that support T cell infiltration and activation within the TME.

### HLA protein expression is inversely correlated with ERS module expression between and within samples

To investigate how ERS activity influences antigen presentation, CyCIF was used to analyze HLA protein expression in cells within 20µm-radius CNs, across the 28 DF/HCC samples. HLA-ABC (MHC class I) and HLA-DPB1 (MHC class II) expression were strongly correlated with each other (SCC = 0.47 **Fig. 6A left**) and positively associated with T cell presence (**Fig. 6A middle and right)**. These data reveal that higher HLA expression correlated with increased T cell aggregation, but this relationship plateaued when T cell counts in a CN reached approximately five, suggesting that antigen presentation supports T cell clustering but alone does not support continuously increased T cell numbers and clusters.

**Figure 6.**
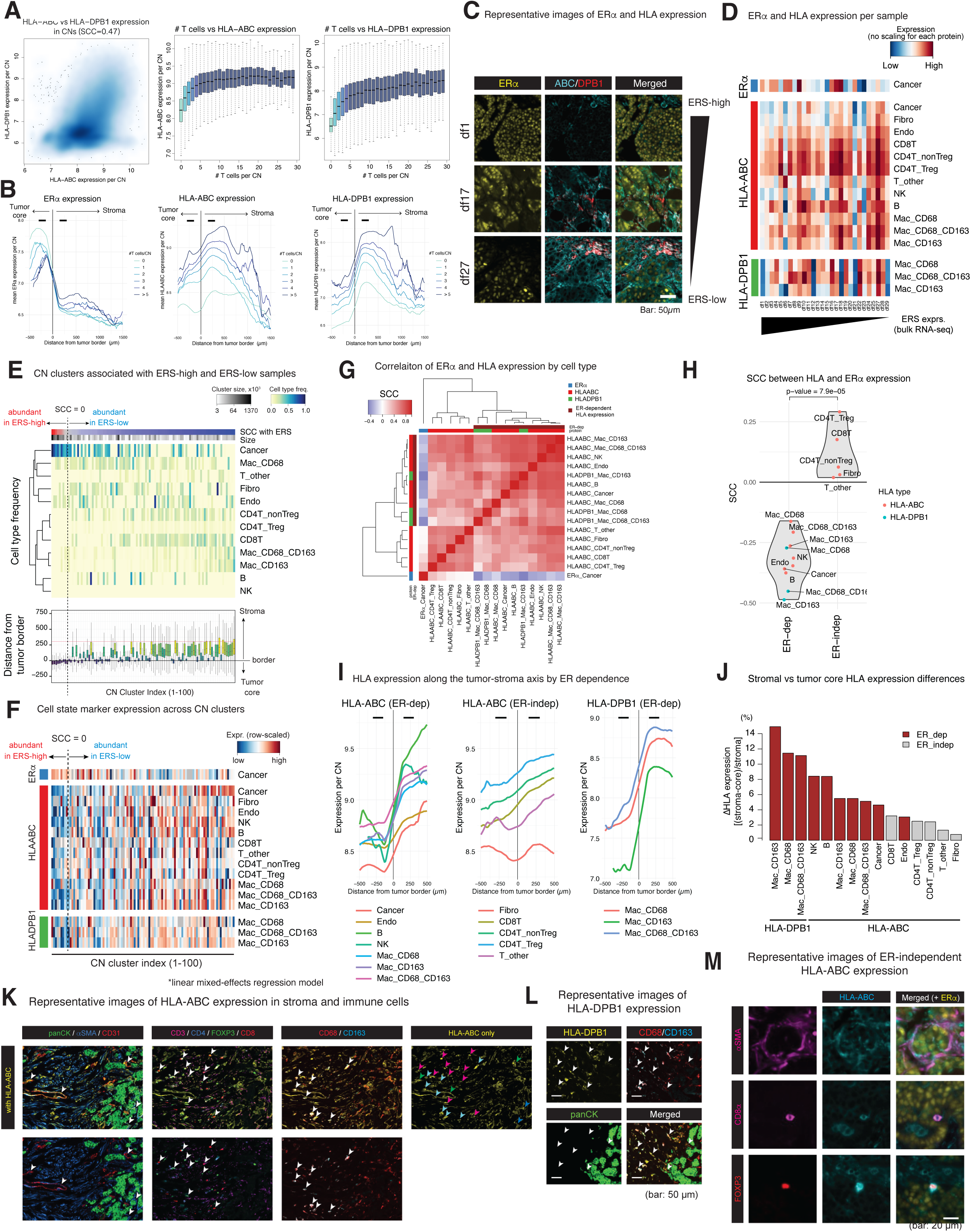
ER-dependent HLA expression is associated with peritumoral T cell infiltration. **A.** Relationship between HLA expression and T cell density in 20µm cellular neighborhoods. Expression of HLA-ABC and HLA-DPB1 in CNs randomly selected from 28 samples (left). HLA-ABC (middle) and HLA-DPB1 (right) expression in CNs stratified by the local T cell density. **B.** Mean protein expression of ERα (left), HLA-ABC (middle), and HLA-DPB1 (right) along the tumor-stroma axis. Curves are stratified by the number of T cells per cellular neighborhood. For HLA expressions, CNs are further grouped by the presence or absence of T cells nearby. Two segments, representing regions within 150-300µm from the tumor border in both tumor core and stroma, were used for comparison of mean protein expression (See **Fig. S10B**). **C.** Representative images of ERα, HLA-ABC, and HLA-DPB1 staining in tumors with low, intermediate, and high ERS levels. The images demonstrate a positive correlation between ERS transcriptional activity and ERα expression in tumor cells, and an inverse correlation with HLA expression. Within each sample, ERα and HLA expression also exhibited an anti-correlated spatial pattern. **D.** Heatmaps depicting cell state marker expression for each cell type, stratified by samples, which shows between-sample diversity of cell state marker expression. **E.** Overview of 100 CN clusters sorted by their prevalence in ERS-high samples, including cluster size, cell type frequency, and distance from the tumor-stromal border. **F.** Heatmap of relative cell state marker expression per cell type among CN clusters. Colors are row-scaled to compare expression across the 100 CN clusters for each cell type. **G.** Spearman correlations between cell state marker expression per cell type across 100 CN clusters. HLA expressions inversely correlated with ER expression in the same CN clusters are highlighted in the top bar, “ER-dep”. **H.** Violin plot of SCC values between ER expression and HLA expressions for each cell type, stratified by ER-dependence. **I.** HLA expression by cell-type along the tumor-stroma axis, grouped by the HLA classes and the ER dependence. ER-dependent cell types show higher HLA expression in stroma over tumor core. **J.** Difference in HLA expression between stroma and the tumor core (ΔHLA) for each HLA class and the cell type. The ΔHLA is normalized by dividing by stromal expression. **K-L.** Exemplar images for HLA-ABC (**K**) and HLA-DPB1 (**L**) expression and multiple cell lineage markerst in tumor samples. **M.** Representative images showing ER-independent HLA-ABC expression in fibroblasts, CD8+ T cells, and Foxp3+ T cells surrounded by ERα-expressing cancer cells. Refer to **Figure S10** for additional details.

Comparison of HLA and estrogen receptor alpha (ERα) protein expression within CNs along the tumor-stroma axis revealed a clear spatial dichotomy: HLA expression was lower in the tumor core, while ERα expression was higher (**Fig. 6B,C**, **Data S1-2**). CNs with more T cells showed increased HLA expression in both the tumor core and stroma, accompanied by lower ERα expression in tumor core, underscoring the link between HLA expression and T cell infiltration (**Fig. S10A**).

Across samples, HLA-ABC and HLA-DPB1 protein expression was inversely correlated with ERS module activity derived from bulk RNA-seq data (**Fig. S10B**). HLA-ABC expression was most prominent in lymphocytes, followed by myeloid cells, stromal cells, and cancer cells. Conversely, HLA-DPB1 expression was predominantly observed in macrophages, especially Mac_CD68_CD163 cells (**Fig. 6D**, **Fig. S10C**). To investigate intra-tumoral variability, CNs were clustered within samples based on cell type frequency profiles. CN clusters dominant in ERS-high samples were primarily composed of cancer cells in the tumor core, while those in ERS-low samples were enriched with T cell and macrophage subsets associated with high APM/TC expression (**Fig. 4C**) and were predominantly located in the stroma. These ERS-low clusters also exhibited higher levels of HLA-ABC and HLA-DPB1 expression (**Fig. 6E**, **Fig. S10D**).

To further explore the relationship between ERα and HLA proteins across the CN clusters, we computed the mean expression levels of ERα, HLA-ABC and HLA-DPB1 for each cell type using a linear mixed-effects model. ERα expression was generally higher in CN clusters prevalent in ERS-high samples, whereas HLA-ABC and HLA-DPB1 expression were elevated in ERS-low clusters (**Fig. 6F**). Correlation analysis revealed that ER-dependent HLA expression, such as in cancer cells and macrophages, was more negatively associated with ERα expression compared to ER-independent HLA expression, such as in T cells and fibroblasts (**Fig. 6G,H**). These distinctions were more pronounced within samples (i.e., intra-tumoral variability) than across samples (i.e., inter-tumoral variability), underscoring the value of within-sample analyses in revealing spatial heterogeneity and local cellular interactions. Clustering protein expression patterns across samples did not reveal a strong negative correlation between ERα and HLA expression (**Fig. S10E**), likely reflecting shared tumor features or technical influences at the sample level. In contrast, within-sample analyses are less affected by these factors, providing a clearer view of biologically meaningful spatial heterogeneity and immune-tumor interactions in HR+ breast cancer.

Given these distinct correlation patterns, HLA expression along the tumor-stroma axis was examined based on whether it was influenced by ER signaling. For cell types where HLA expression exhibited ER dependence – including cancer cells, endothelial cells, macrophages (Mac_CD68, Mac_CD163, Mac_CD68_CD163), NK cells, and B cells – HLA-ABC expression was lower in the tumor core compared to the stroma (**Fig. 6I**, **left**). In contrast, for cell types where HLA expression appeared ER-independent, such as T cells and fibroblasts, HLA-ABC expression was more consistent across the tumor core and stroma (**Fig. 6I**, **middle**). HLA-DPB1 expression in macrophages followed an ER-dependent pattern, with higher levels in the stroma compared to the tumor core (**Fig. 6I**, **right**). The differences in HLA expression between the tumor core and stroma may suggest distinct mechanisms of HLA expression based on ER expression (**Fig. 6J**).

These spatial differences suggest that ER signaling selectively modulates HLA expression in specific cell types, influencing antigen presentation capabilities and shaping immune cell dynamics within the TME. Micrographs corroborated these findings, showing cell type-specific patterns of HLA expression in the stroma (**Fig. 6K,L**).In particular, we visually confirmed that fibroblasts and T cells show HLA-ABC molecules in an ER-independent manner, while surrounded by the ER+ tumor cells.

### Endocrine therapy induces changes in ERS and immune module activities in HR+ breast cancer

Our analysis consistently demonstrated an inverse correlation between ERS and immune signaling in HR+ breast cancer, motivating an investigation into how these pathways respond to modulations in ERS activity. To explore this, we analyzed a microarray dataset from the ACOSOG Z1031B clinical trial, which provides pre-and post-treatment data for 214 evaluable HR+/HER2-patients treated with preoperative aromatase inhibitor (AI) therapy in patients with stage II or III ER+ breast cancer (*47, 48*).

Among these patients, 34 exhibited increased Ki67 levels post-treatment, indicating AI resistance, while 180 showed decreased Ki67, suggesting AI sensitivity. At baseline, AI-sensitive tumors displayed higher ERS module expression and lower TNFα/NF-κB and APM/TC module activity, with less pronounced difference in the IFN-I module (**Fig. 7A**, **Fig. S11A**). Post-treatment, responsive tumors showed decreased ERS module expression and increased activity of TNFα/NF-κB and APM/TC modules, while IFN-I remained relatively unchanged (**Fig. 7B-C**). In contrast, AI-resistant tumors exhibited decreased ERS activity with minimal changes in immune module activity, suggesting that in AI-sensitive tumors, ERS suppression is specifically linked to TNFα/NF-κB activation and increased immune activity (**Fig. 1E,H**) (*72*).

**Figure 7.**
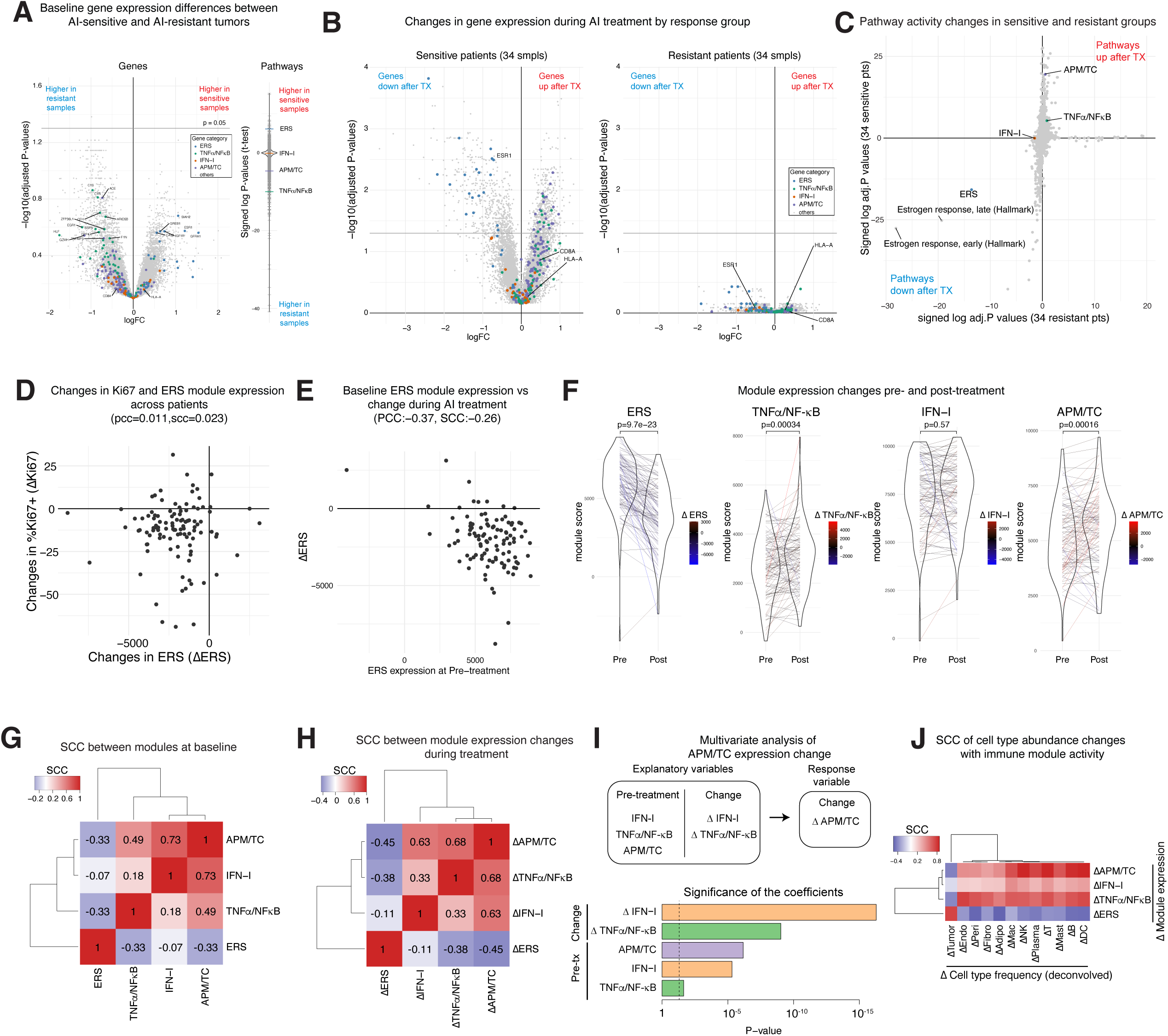
TNFα/NF-κB activation linked to ER signaling suppression in AI-responsive HR+ breast cancer. Analysis of ACOSOG Z1031B trial samples. **A.** Differential gene expression analysis between AI-sensitive and AI-resistant patients at pre-treatment. Volcano plot showing gene expression differences between 34 AI-resistant and 34 AI-sensitive patients (left). Violin plot illustrating pathway activity differences between the two groups (right). **B.** Volcano plots of genes altered by AI therapy in sensitive (left) and resistant (right) patient groups. **C.** Changes in pathway activity during AI treatment in sensitive (x-axis) and resistant (y-axis) patients. Pathways in the bottom-left quadrant (third quadrant) are those downregulated after AI treatment in both groups. **D.** Relationship between changes in Ki67 positivity and ERS module expression across all 214 patients. **E.** Relationship between ERS module expression at pre-treatment and the change in ERS activity during treatment. **F.** Expression of the four modules before and after the treatment. Matched patients at two time points are connected by lines, with colors indicating the direction of change. **G.** SCC between the four modules at pre-treatment. **H.** SCC between changes in module expression during AI treatment. **I.** Multivariate linear regression analysis identifying factors contributing to change in APM/TC module expression. Explanatory variables include immune module expression at the pre-treatment and their changes during treatment. **J.** Correlation between changes in module expression and cell type abundance during AI treatment. Refer to **Figure S11** for additional details.

Although changes in Ki67-positive cell frequency (ΔKi67) was the primary measure of therapeutic response in the trial, changes in ERS module expression (ΔERS) were not correlated with ΔKi67 (**Fig. 7D**). However, most tumors showed a consistent pattern of decreased ERS, increased TNFα/NF-κB and APM/TC activity, and negligible changes in IFN-I (**Fig. 7F,G**). Notably, changes in APM/TC module expression (ΔAPM/TC) correlated more strongly with changes in TNFα/NF-κB (ΔTNFα/NF-κB) than IFN-I (ΔIFN-I), reflecting trends observed in ΔKi67-based patient stratification (**Fig. 7H**). These changes were inversely correlated with their baseline expression levels, indicating that tumors with higher baseline ERS and lower immune module expression exhibited a greater capacity for dynamic response to therapy (**Fig. 7E**, **Fig. S11B**).

Multivariate analysis identified ΔIFN-I and ΔTNFα/NF-κB as the strongest predictors of ΔAPM/TC, with pre-treatment immune module expression also contributing (**Fig. 7I**). Cell type abundance analysis confirmed expected associations between module activity and specific cell types: ERS with cancer cells and APM/TC with lymphocytes (**Fig. 7J**, **Fig. S11C-E**). Notably, TNFα/NF-κB module expression correlated most strongly with endothelial cell abundance, suggesting that endothelial cells may play a key role in modulating this pathway and enhancing immune infiltration. These findings support the hypothesis that AI therapy suppresses ERS, releasing TNFα/NF-κB suppression, and facilitating APM/TC activation. TNFα/NF-κB activity appears to be partially driven by endothelial cells, creating a more permissive environment for immune infiltration.

### Summary of ER Signaling and Immune Exclusion Mechanisms in HR+ Breast Cancer

Here we present robust evidence that ER signaling influences immune and stromal dynamics in HR+ breast cancer (**Fig. 8**). In ERS-high tumors, heightened ER signaling suppresses both IFN-I and TNFα/NF-κB responses, leading to reduced non-cancer cell infiltration, diminished MHC expression, and limited PD-L1 upregulation and angiogenesis. These effects collectively contribute to a lack of stromal T cells in the TME. By contrast, ERS-low tumors exhibit increased IFN-I and TNFα/NF-κB activity, which may drive non-cancer cell recruitment, enhance MHC expression, and promote angiogenesis and PD-L1 expression. These changes are associated with an increased presence of stromal T cells, which predominantly express PD-1, suggesting potential ability to stimulate with anti-PD-1 or anti-PD-L1 therapy. The distinct regulation of IFN-I and TNFα/NF-κB pathways by ER signaling provides mechanistic insight into immune exclusion in HR+ breast cancer and highlights pathways that may be targeted to improve therapeutic outcomes.

**Figure 8.**
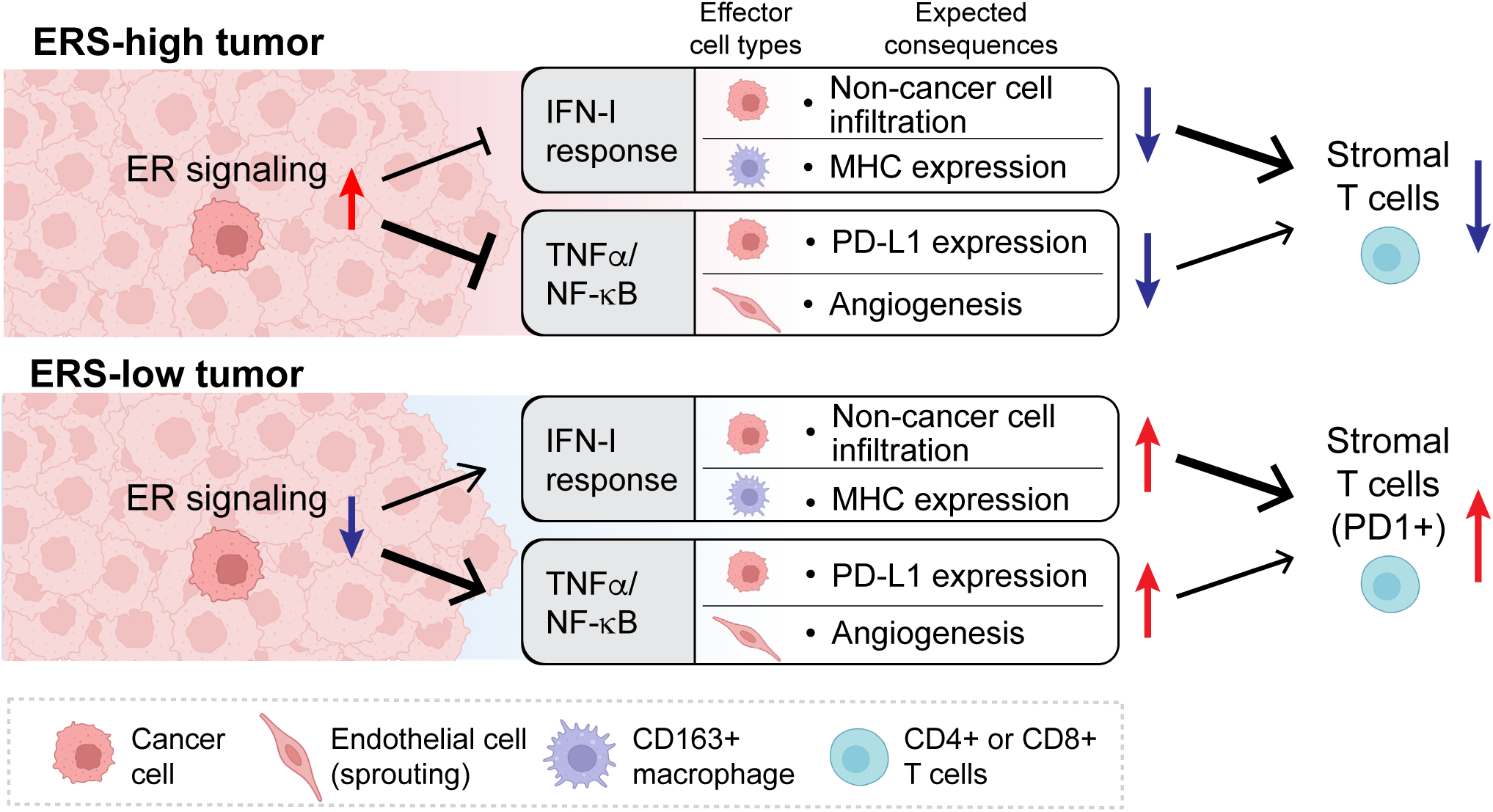
Updated model illustrating ER signaling-induced T cell exclusion in HR+ breast cancers.

## Discussion

Understanding the immunology of HR+ breast cancer is essential for identifying patients most likely to benefit from immunotherapy strategies, as adding immunotherapy to an unsuitable population may not improve efficacy but may add additional toxicity. HR+/HER2-breast cancer is considered immunologically cold due to its low levels of PD-L1 expression, low T cell abundance, and low tumor mutational burden. Despite these challenges, we found that HR+ breast cancers displayed significant variability in the expression of ER signaling related genes, T cell abundance and immune landscape. Our findings corroborate prior studies that highlight an inverse relationship between ER signaling and immune activation in HR+ breast cancer (*45*). By combining transcriptomic and spatial analyses, we identified associations between ERS and immune activity, including reduced recruitment of non-cancer cells to the TME and suppression of antigen presentation. These findings provide a foundation for exploring how heightened ERS suppresses immunogenicity and impedes T cell infiltration (**Fig. 8**).

Using TCGA and METABRIC datasets, we identified distinct gene modules linked to ER and immune signaling. The ERS module, which captures both initial and sustained estrogen responses, was predominantly expressed in cancer cells, while immune-related modules were expressed in non-cancer cell types. Specifically, TNFα/NF-κB was expressed in fibroblasts and endothelial cells, IFN-I signaling was most prominent in macrophages, and APM/TC, associated with antigen presentation and T cell function, was expressed in various immune cells. Importantly, analysis of CCLE breast cancer cell lines revealed cancer cell-intrinsic inflammatory pathways, including TNFα/NF-κB and IFN-I signaling, within cancer cells. Notably, the TNFα/NF-κB module included PD-L1, suggesting that ERS directly suppresses PD-L1 expression in cancer cells (*73*). Similarly, IFN-I expression in cancer cells was linked to antigen-presentation machinery, indicating a direct role for type I IFN signaling in modulating immune responses (*74, 75*).

Our findings emphasize the cancer cell-intrinsic role of IFN-I signaling in maintaining immune cell recruitment to the TME. Suppression of cancer cell-intrinsic IFN-I signaling by ERS correlates with reduced antigen presentation and T cell exclusion, unveiling a crucial mechanism by which ER signaling may drive immune evasion in HR+ breast cancer. Genes involved in antigen processing and presentation, such as HLA-A, B2M, PSMB9 and NLRC5, are expressed within the intrinsic IFN-I module. These genes, along with the type I IFN response, were not only associated with T cell abundance, but also linked to various stromal and immune cell recruitment. This underscores the central role of the cancer cell-intrinsic type I IFN signaling in sustaining immune cell presence, a process that ERS may actively suppress.

Spatial analyses further elucidated the interplay between ERS, immune modules, and cellular composition. CyCIF analysis of 29 primary HR+ breast cancer samples revealed that CD4+ and CD8+ T cells were enriched in ERS-low tumors and co-localized with macrophage subsets (CD163+ and CD68+/CD163+) in the stroma. TNFα/NF-κB and IFN-I activity were strongly associated with endothelial cells and stromal macrophages, respectively, indicating that these immune pathways may shape the spatial immune landscape. While T cells formed clusters near the tumor border in ERS-low samples, these cells predominantly expressed PD-1, indicating exhaustion. PD-1+ T cells were closely associated with CD163+ macrophages in the stroma, of ERS-low samples, suggesting a mechanism of suppression and explaining the limited efficacy of PD-1/PD-L1 blockade therapy, despite the presence of PD-1+ T cells in this context. Although CyCIF did not include markers specific to dendritic cells (DCs), single-cell and single-nucleus transcriptomic data from the WEHI and DF/BCC cohorts suggested that DCs co-occur with macrophages and T cells. Given their critical role in antigen presentation and immune activation, this raises the possibility that DCs contribute to the observed immune modulation in HR+ breast cancer and warrants further investigation to elucidate their spatial distribution and function.

In addition to ER signaling, several mechanisms of T cell exclusion have been proposed, including TGFβ signaling and the role of cancer-associated fibroblasts (CAFs). CAFs are known to remodel the extracellular matrix and secrete immunosuppressive factors such as TGFβ, creating barriers to T cell infiltration (*76–78*). However, in our analysis of HR+ breast cancer, extensive CAF presence was not a common feature across most samples, suggesting CAFs may play a more limited role in T cell exclusion within the DF/BCC cohort. Instead, our findings highlight ER signaling as a primary mechanism of immune exclusion, particularly through suppression of IFN-I and TNFα/NF-κB pathways. These results suggest that HR+ breast cancer employs unique strategies for immune evasion, distinct from those in other solid tumors, such as pancreatic cancer.

Insights from the ACOSOG Z1031B trial provide additional support for the role of ER signaling in immune modulation. Endocrine therapy with aromatase inhibitors (AIs) reduced ERS activity, leading to increased TNFα/NF-κB and APM/TC module activity in AI-sensitive tumors. Although IFN-I activation was less pronounced, multivariate regression analysis confirmed its contribution to APM/TC activation. These findings emphasize the necessity of activating both TNFα/NF-κB and IFN-I pathways to induce robust T cell infiltration and activation in HR+ breast cancer. Therapeutic strategies that combine ER inhibition with modulators of TNFα/NF-κB and IFN-I signaling could enhance the efficacy of immunotherapy.

Clinical trials provide some suggestive evidence for this approach. For example, the DOLAF trial evaluated the combination of the PARP inhibitor olaparib, a selective ER degrader fulvestrant, and the PD-L1 inhibitor durvalumab, in metastatic HR+ breast cancer (*79*). Olaparib has been shown to induce cGAS-STING signaling and IFN-I responses in patients with *BRCA-*associated breast cancer (*80, 81*). While the trial was not randomized, and included patients beyond those with *BRCA* mutations, the triple combination revealed meaningful PFS and raises the possibility that combining TNFα/NF-κB and IFN-I activation with inhibition of ER signaling could improve outcomes by enhancing T cell infiltration and activation. Importantly, taken together, our work exemplifies how targeting multiple pathways may overcome immune exclusion and improve therapeutic responses.

### Limitations of this study

While the correlations between ERS and immune signaling observed in this study were validated across multiple datasets, they should be interpreted with caution as suggestive rather than conclusive evidence of causality. Additionally, our analyses primarily focused on untreated baseline samples. Further studies examining on-treatment samples, such as those from the ACOSOG Z1031B trial, are warranted to validate the dynamics of module expression and their spatial distribution in response to therapy and how these changes correlate with patient outcomes. Expanding these investigations could provide a more comprehensive understanding of how ERS and immune signaling interplay to shape therapeutic responses.

## Acknowledgements

We are grateful for expertise and help from the following core facilities: The Brigham and Women’s Center for Advanced Molecular Diagnostics Research Core Lab and The Multiplex Imaging Platform of the Laboratory of Systems Pharmacology (LSP) at Harvard Medical School. We are grateful for the contributions of Dr. Isabelle Bedrosian for sharing 74 breast cancer samples, collected at MD Anderson Cancer Center.

## Author contributions

Conceptualization: KS, YXC, JG, JA, EAM, JLG

Data curation: KS, DM, YXC, JG, ZJ, LK

Formal analysis: KS, DM

Funding acquisition: EAM,

JLG Investigation: DM, ZJ

Methodology: KS

Project administration: KS, JLG

Software: KS, DM, ZJ, JH, JM, NH, RK,

Resources: DM, ZJ

Supervision: KS, EAM, JLG

Validation: KS

Visualization: KS

Writing – original draft: KS, JLG

Writing – review & editing: DM, YXC, KZ, JG, ZJ, SJS, RP, LK, JH, JM, NH, RK, AG, AN, CW, GA, SM, SMT, AGW, RJ, PKS, JA, EAM

## Data and materials availability

all curated code is available at https://github.com/GuerrieroLab/HR-APM-Paper. Processed transcriptome and CyCIF data from DF/BCC cohort will be made accessible upon publication in a peer-reviewed scientific journal.

