## Supplementary material for "An estrogen receptor signaling transcriptional program linked to immune evasion in human hormone receptor-positive breast cancer": Materials and Methods and Supplemental Figure Legends

#### Manual curation of the gene panels related to ERS and immune activity

Gene panels for ERS and immune activity, specifically focusing on antigen presentation machinery and T cell signaling, were manually curated from various sources, including MSigDB, STRING, and literature (Andruska et al., 2015; Bhardwaj et al., 2019; Chaudhary et al., 2017; Dutertre et al., 2010; Manavathi et al., 2014; Shanmugam et al., 1999; Vella et al., 2020; Yager and Davidson, 2006). Of the 988 genes identified, 778 were measured in both the TCGA and METABRIC cohorts and included in the study for further analysis (**Table S1**).

#### Data collection

All datasets except DF/BCC were collected electronically. Bulk transcriptome data and clinical metadata for TCGA, METABRIC, and CCLE were retrieved from cBioPortal (https://www.cbioportal.org/). Transcriptome data for 74 breast cancer patients from MD Anderson Cancer Center were provided by Dr. Bedrosian. Bulk RNA-seq from the ACOSOG Z1031B trial and scRNA-seq data from the WEHI cohort were downloaded from GEO (Accession #GSE87411 and #GSE161529). DF/BCC cohort data were collected as described below.

#### Correlation analysis

Correlation analyses employed Pearson's correlation coefficient (PCC) for sample sizes ≤40, levering its staitiscal power, and Spearman's rank correlation coefficient (SCC) for larger samples (>40) to account for potential non-linear relationships.

#### Preprocessing for TCGA and METABRIC transcriptome datasets

For TCGA (719 patients) and METABRIC (1,904 patients), gene expression values were log2-transformed, and breast cancer subtypes categorized by HR and HER2 status via immunohistochemistry (IHC). Duplicates were removed by retaining the most abundantly expressed gene variant to create a clean dataset.

Comparisons of gene expression between TCGA and METABRIC identified a discrepancy in expression levels, attributed to the reduced sensitivity of the METABRIC microarray platform. In METABRIC, a histogram of expression levels for individual genes across patients peaked near zero, indicating that many genes were not expressed or were lowly expressed but could not be distinguished from background noise. To address this, a signal threshold based on the 99.9th percentile of the background distribution was set in METABRIC. Mean expression for 56% of all genes fell below this threshold, suggesting that a substantial portion of gene expression was affected by noise. In contrast, TCGA did not exhibit a similar peak at low expression levels, indicating a higher sensitivity in detecting expressed genes.

We then examined gene co-expression patterns by comparing the Spearman correlation coefficients (SCC) of 778 genes between the datasets. The SCC values were consistently higher in TCGA, indicating a better signal-to-noise ratio in this dataset. Genes flagged as artifacts in METABRIC were found to have substantially lower correlations with other genes, further supporting the idea that low-expressed genes in METABRIC may be influenced by noise. Consequently, we used the TCGA dataset for primary analyses, relying on METABRIC data for validation.

As part of the METABRIC preprocessing, an artifact removal step was applied to specific genes. For instance, *HLA-A* exhibited undetectable expression in a large subset of samples, which was likely due to experimental biases. To refine artifact detection, we applied a model selection process based on the Bayesian Information Criterion (BIC) using mclustBIC in mclust R package, which identified 20 genes, including *HLA-A*, that were likely affected by technical artifacts and showed no expression in certain patient samples. Expression values for these genes that were indistinguishable from the background were set to missing (NA) in samples where their expression fell below the threshold.

All gene expression data were standardized by z-score transformation across all samples for downstream analyses.

**Co-expressing gene module identification**

To identify co-expressing gene modules, we calculated pairwise distances between 778 genes from our manually curated list, using 1 minus the Spearman correlation coefficient (SCC) as the distance metric. We applied hierarchical clustering with average linkage across hormone receptor-positive (HR+) breast cancer samples in both the TCGA and METABRIC datasets. This method grouped genes into clusters based on their expression patterns, and we inspected these clusters for hallmark genes such as *ESR1*, *CD8A*, and *HLA-A*, which guided the identification of biologically relevant modules.

In the TCGA dataset, we initially identified clusters corresponding to estrogen receptor signaling (ERS), cell cycle (CC), and pan-immune-related genes. The ERS and CC clusters were defined based on their specific gene lists, while the pan-immune clusters were derived from immune-related genes. Hierarchical clustering results were manually refined using the cutree function to select tightly clustered genes, ensuring that only genes with strong co-expression relationships were included in each module.

We repeated the same hierarchical clustering process in the METABRIC dataset to validate and refine these modules. Genes that consistently clustered together in both datasets were included in the final modules. To further enhance module specificity, we refined the gene modules by retaining only those genes that exhibited strong positive correlations within their respective modules in the TCGA dataset. This refinement process led to the exclusion of six genes from the pan-immune modules due to weak correlations, while all genes in the ERS module were retained.

**Gene module characterization with pathway analysis**

Pathway enrichment analysis was conducted using MSigDB Hallmark gene sets to characterize the biological relevance of the ERS, CC, and pan-immune modules. Fisher's exact test was applied to identify pathways enriched within each module, with multiple testing correction performed using the q-value method. Enrichment was defined as q-value < 1e-5, and gene modules were characterized by their unique pathway enrichments.

**Breast cancer subtype analysis in TCGA and METABRIC cohorts**

Correlation matrices for gene modules (ERS, TNFα/NF-κB, IFN-I, APM/TC) were computed for four breast cancer subtypes: HR+/HER2-, HR+/HER2+, HR-/HER2-, and HR-/HER2+ in both the TCGA and METABRIC datasets. Heatmaps and violin plots were generated to visualize and compare module correlations within and between subtypes. Since *ESR1* gene expression or ERS module activity clearly differentiated HR+ from HR- samples (AUROC = 0.97 for *ESR1* gene and 0.96 for ERS module), the relationship between ERS and immune-related modules was further explored by analyzing their activity in individual samples pooled across all subtypes. GSVA was performed on TCGA and METABRIC expression data to quantify pathway activity across subtypes, with Z-score normalization used to compare the enrichment scores for the modules across all samples.

**Validation with MD Anderson Cancer Center cohort data**

Gene module co-expression patterns observed in TCGA and METABRIC datasets were validated using the MD Anderson Cancer Center (MDACC) cohort, consisting of RNA-seq data from 74 tumors. Spearman correlation coefficients between module genes were calculated for each breast cancer subtype, and the mean correlation within and between gene modules was visualized.

**Preprocessing of scRNA-seq data from the WEHI cohort and cell type identification**

Single-cell RNA-seq data from 34 patients in the WEHI cohort was processed to uncover distinct cell populations associated with various breast cancer subtypes. Subtype classifications (HR+, HER2+, TNBC) were provided by the data contributors, and these classifications were further validated using a custom basality score that assesses subtype-specific keratin expression (e.g., luminal markers like KRT8, KRT18, and basal markers like KRT5, KRT17).

To comprehensively explore immune and stromal populations, we applied the Louvain clustering algorithm in Seurat's FindClusters function and marker-based identification via FindAllMarkers function. This dual approach allowed for the identification of broad cell categories, including myeloid, lymphoid, and epithelial cells. Further refinement within these groups led to the identification of subtypes such as macrophages and dendritic cells among myeloid cells, and distinct T cells, B cells, plasma cells, and NK cells within lymphocytes, based on differential gene expression profiles. These classifications was visually confirmed using UMAP plots, which showcased the expression of key marker genes, ensuring the precise identification of each cell type.

**Analysis of gene expression in the WEHI cohort**

Gene module activity in the WEHI cohort was assessed using single-sample GSEA (ssGSEA) via the GSVA package. Enrichment scores were computed for each module and visualized on UMAP plots to illustrate the spatial distribution of module activity across different cell types.

For a more granular analysis of module activity, we calculated the average expression of each module per sample and per cell type. A matrix deconvolution method was applied to partition the overall module scores into contributions from individual patient samples and distinct cell types. Additionally, gene-level analysis across all modules confirmed that genes exhibited consistent expression patterns across cell types and patient samples within the same module, while demonstrating distinct patterns between different modules. This gene-level analysis underscored the co-regulation of genes within each module and complemented the module-level analysis by providing deeper insight into the gene-specific contributions to overall module activity.

Finally, cell type frequencies were computed and correlated with module activities, exploring how the abundance of different cell types influences gene module expression across patient samples in the breast cancer microenvironment.

**Identification of genes expressed in CCLE Cell Lines**

PAM50 subtypes of 54 breast cancer cell lines from CCLE were taken from Jiang et al.(*1*), including 19 luminal A or B subtypes representing HR+ breast cancer. Gene module analyses were conducted using both the subset of 19 luminal lines and the entire set of 54 cell lines, enabling comparisons within HR+ subtypes and across broader breast cancer subtypes.

To analyze gene expression in the CCLE cell lines, we compared the expression of genes across breast cancer cell lines, distinguishing between expressed and non-expressed genes. Genes highly expressed in T and NK cells from the WEHI dataset were absent in the CCLE cell lines, which we further explored using the ROCR package. This analysis established a threshold to differentiate between gene expression patterns typical of tumor cells versus T/NK cells.

Next, gene expression correlations were examined in both the full dataset (54 lines) using Spearman correlations and the HR+ subset (19 lines) using Pearson correlations. Heatmaps and hierarchical clustering were employed to visualize co-expression patterns, revealing conserved gene modules across the datasets.

The preserved genes were labeled with their original module names (e.g., IFN-I), with the addition of "int" or "intrinsic" to denote cancer cell-intrinsic modules. These intrinsic modules were further divided into submodules based on the original whole-tumor module they derived from, such as "IFN-sub" or "APM-sub."

Finally, pathway enrichment analysis using Fisher's exact test was conducted to identify pathways enriched in the preserved gene sets, revealing their functional relevance across breast cancer subtypes.

**Collection of surgical specimens from the DF/BCC cohort**

Transcriptome and Cyclic Immunofluorescence (CyCIF) data were obtained from 29 primary treatment-naïve HR+ breast cancer surgical resections, collected from patients under Dana-Farber Cancer Institute protocol # 93-085. Fresh tissues were split in half with a sterile razor and either placed in a cryovial for immediate flash freezing in liquid nitrogen or placed in 10% neutral buffered formalin for fixation for 24 hours prior to paraffin embedding. Estrogen receptor, progesterone receptor, and HER2 expression were confirmed by immunohistochemistry (IHC). Tissues with Equivocal (2+) HER2 IHC staining were tested for *ERBB2* amplification by fluorescent in-situ hybridization (FISH).

**RNA extraction and sequencing for bulk RNA-seq**

RNA was extracted from FFPE sections, quantified, and sequenced using 2x150 bp reads on an Illumina NovaSeq sequencer. Sequencing reads were aligned to the GRCh38 reference genome using STAR2.7.2b. The raw counts were normalized by Transcripts Per Million (TPM) method. The ensemble IDs were converted to gene symbols using Gencode Human Release 44.

**Single-nucleus GEM generation and gene expression library construction**

Nuclei isolation was performed as previously described (*2*). Low-retention microcentrifuge tubes (Fisher Scientific, Hampton, NH, USA) were used throughout the procedure to minimize nuclei loss. Briefly, tissues were manually dissociated by chopping with fine spring scissors for 10 minutes, homogenized in TST solution, filtered through a 30-µm MACS SmartStrainer (Miltenyi Biotec, Germany), and pelleted by centrifugation for ten minutes at 500g at 4°C. Nuclei pellets were washed with 1 mL of resuspension buffer (1X PBS + 1% BSA) and filtered again, centrifuged at 500g for 5 minutes at 4°C and then resuspended in 100 µL of resuspension buffer and trypan blue-stained nuclei were counted manually using INCYTO C-Chip Neubauer

Improved Disposable Hemacytometers (VWR International Ltd., Radnor, PA, USA) under a brightfield microscope. This process was performed for each sample independently. The yield and quality of nuclei were adequate to process further for single-nuclei RNA sequencing. A droplet-based microfluidic Chromium Controller (10X Genomics, Inc., Pleasanton, CA, USA) was used to generate single-nuclei Gel Beads-in-emulsions (GEMs), as per the Chromium Single Cell 5' Reagent Kits (Cat no. 1000263) User Guide RevE (v2 Chemistry Dual Index; 10X Genomics demonstrated protocol). Approximately 10,000 nuclei were loaded with 10X-barcoded gel beads and partitioning oil including RT master mix for GEM generation. Next, GEMs were incubated to produce 10x barcoded, full-length double-stranded cDNA from polyadenylated mRNA. The Agilent High Sensitivity DNA Kit assessed the resulting cDNA with the Agilent Bioanalyzer 2100 (Agilent Technologies, Lexington, MA, USA). Approximately 50 ng of cDNA were processed for gene expression library construction. and libraries were also quantified on the Agilent Bioanalyzer 2100. Next, libraries were assessed on the Agilent Bioanalyzer 2100. All the libraries were normalized and pooled for sequencing on Illumina NovaSeq platforms (Illumina, Inc., San Diego, CA, USA).

**Single-nucleus RNA-seq data pre-processing for DF/BCC data**

Pooled samples were demultiplexed using 10x Cellranger mkfastq. Read pairs were aligned to the hg38 reference sequence using a 10x Cellranger count, which utilizes STAR (*3*). Cellbender remove-background (*4*) was run on aligned, barcoded reads to distinguish cell-containing droplets from empty droplets and retrieve background-free gene expression profiles. Cellbender-filtered matrices were analyzed in RStudio using Seurat v5(Hao, et al., 2023). Here, data was further filtered by retaining cells with $\geq$ 200 detected genes, and genes if they were expressed in $\geq$ 3 cells. Cells with < 20% mitochondrial read were retained. All samples were merged into a single Seurat object and cells were integrated using Harmony (*5*). Dimensionality reduction was performed using the Uniform Manifold Approximation and Projection (UMAP) algorithm from matplotlib to cluster cells and preserve similarities in the original gene expression space (*6*). Individual clusters were annotated using canonical marker expression determined by the FindAllMarkers function within Seurat v5.

**Bulk and single-nucleus RNA-seq analysis of DF/BCC cohort**

For the bulk and snRNA-seq analyses of the DF/BCC cohort, we applied the same pipeline as used for the bulk data from TCGA/METABRIC and for the single-cell data from the WEHI cohort.

**CyCIF staining and data pre-processing**

CyCIF was performed on FFPE sections with antigen retrieval, blocking, and repeated staining, imaging, and bleaching cycles as previously described (*7*). Briefly, FFPE slides were deparaffinized and underwent antigen retrieval using a LEICA Bond Rx. Slides were blocked overnight at 4°C in Super Block Blocking Buffer (Thermo; 37515) containing secondary F(ab')2 fragments against each species of primary antibody used. Slides were bleached using $NaOH/H_2O_2$ solution and washed with PBS prior to staining. Slides were stained with primary antibodies for each cycle overnight at 4°C and were washed the following day with PBS. DNA was visualized using 1ug/mL Hoechst 33342 (Thermo; 62249). Coverslips were mounted with 70% glycerol and imaged on the CyteFinder II automated slide scanning instrument (RareCyte) at 20x magnification. Primary antibodies that are used for the analysis are as follows: HLA-ABC

(Abcam; ab70328), CD3 (Abcam; ab11089), CD31 (Abcam; ab218582), CD4 (R&D Systems; AF-379-NA), CD68 (Cell Signaling; 24850), CD45 (Biolegend; 304060), HLA-DBP1 (Abcam; ab201347), aSMA (R&D Systems; IC1420S), Granzyme B (Abcam; 225472), FOXP3 (Thermo; 41-4777-82), CD8A (Thermo; 50-0008-82), panCK (Novus; NBP2-33200AF750), CD163 (Abcam; ab218293), CD57 (Biolegend; 359612), CD20 (Thermo; 50-0202-80), ERa (Novus; NBP2-33321AF750), pTBK1 (Cell Signaling; 14586S), ISG15 (Santa Cruz; sc-166755 AF594), PD1 (Abcam; ab201825).

In CyCIF analysis, whole slide images from multiple cycles are overlaid, and cells are segmented using the MCMICRO computational platform (*8*). These segmented images are visualized through OMERO (*9*), and specific regions of interest (ROIs) are selected, excluding areas with tissue loss, folding, or imaging artifacts. For each sample, gates are defined using Gater (https://github.com/labsyspharm/gater) to determine whether particular cell lineage markers are expressed, based on fluorescence intensity within the ROIs.

To streamline and extend these analyses, the CycifAnalyzeR package organizes each sample's data into a *Cycif* object, which stores spatial and expression information of segmented cells. These individual *Cycif* objects are then compiled into a *CycifStack* object, enabling batch processing and comprehensive spatial analysis across multiple samples. The package provides tools for various spatial analyses, including computing distances between cells, identifying cellular neighborhoods, and evaluating spatial relationships, such as co-localization of cell types and clustering patterns around tumor and stroma regions. After importing the data and applying the gates, cell types are identified using the defineCellType() function, which applies a hierarchical cell type definition scheme, as shown in Fig. S8.

### Integrative analysis of cancer cell signaling in WEHI and DF/BCC

This analysis examined the relationship between gene expression, cancer cell frequency, and pathway activity across the WEHI and DF/BCC cohorts. First, genes commonly measured in both the scRNA-seq data from WEHI and bulk/snRNA-seq data from DF/BCC were identified. The gene list was refined by filtering for genes where cancer cell expression in individual samples exceeded 10% of the highest expression in other cell types. This ensured the focus on genes predominantly expressed within cancer cells for both datasets.

Cell type frequencies were determined using scRNA-seq for WEHI and CyCIF for DF/BCC, and mean gene expression within cancer cells was calculated for each sample in both cohorts. Correlation analyses were conducted to assess the relationship between gene expression and cancer cell frequency across datasets. Pathway activity was computed using fgsea, and Pearson correlations were calculated between cancer cell frequency and pathway activity in cancer cells across cohorts. Pathways consistently correlated with cancer cell frequency were identified, providing insights into cancer cell-intrinsic factors.

Additionally, correlation and pathway enrichment analyses were performed using bulk RNA-seq data from the DF/BCC cohort to further investigate the relationship between cancer cell frequency and pathway activity in whole tumors (compared to cancer cells).

### Spatial analysis of CyCIF data

Spatial analyses were conducted using the CycifAnalyzeR package (Version 1.0) to explore cell interactions within tissue architecture. The analysis focused on investigating tumor-stroma boundaries, the spatial distribution of cell types along the tumor-stroma axis, and calculating cell neighborhood statistics. Additionally, the spatial organization of immune and non-immune cells was examined, providing insights into tissue structure and cell communication.

**Tumor-stroma ratio and cell type frequency analysis**

The spatial analysis began by examining the distribution of cell types across the tumor-stroma axis. Using the defineTumorStroma() function, cells were classified into tumor or stroma regions based on their distance from the tumor boundary, which was divided into 20-micron intervals (bins). The tumor-to-stroma ratio for each cell type was then calculated, normalizing for the total number of non-cancer cells in each sample. This analysis provided insights into the abundance of different cell types within the tumor core and stroma. The relative frequency of each non-cancer cell type along the tumor-stroma axis was computed and smoothed using a 10-bin moving average, revealing trends in their spatial distribution.

**Endothelial cell network size and spatial distribution**

Endothelial cells were identified based on CD31 expression, and the DBSCAN algorithm was used to cluster cells into vascular networks. The size of these networks was calculated and plotted as a function of the distance from the tumor boundary, with a moving average applied to smooth fluctuations and highlight consistent trends in endothelial organization along the tumor-stroma axis.

**Computing local density and protein expression through cellular neighborhoods**

The computeCN() function was used to calculate the local density of each cell type by counting corresponding cells within a 20-micron radius. This approach enabled the analysis of spatial co-localization and interactions among specific cell types within the tumor microenvironment. For protein expression, two levels of analysis were performed: aggregate expression, measuring the mean protein expression across all cells within CNs, and cell type-specific expression, focusing on mean protein expression for individual cell types. Protein markers such as HLA-ABC, HLA-DPB1, ERα, and PD1 were examined to assess their activity across immune and stromal populations.

**CN Clustering**

To capture variations in cellular neighborhoods, up to 10,000 CNs were sampled per patient and pooled for analysis. CNs with at least 10 cells were included, and PCA was applied to cell type frequency data to reduce dimensionality. K-means clustering was used to group CNs into 100 clusters based on cell type composition and protein expression. The resulting clusters were analyzed for their prevalence in ERS-high samples and examined through correlation analyses to understand the organization of cells and proteins in the tumor microenvironment.

**Within-sample protein expression using linear mixed-effects models**

To quantify protein expression by cell type and CN cluster, linear mixed-effects models were applied: $Expression \sim CN\ cluster + (1|Sample)$

This model captures cluster-specific protein expression patterns while controlling for sample-to-sample variability. The resulting coefficients revealed how protein expression varied spatially, providing insights into the influence of local microenvironments on immune and stromal cell populations, particularly regarding ERS signaling and immune checkpoint regulation.

**Assessment of ER-dependent and independent HLA expression**

To assess the relationship between T cells and HLA expression within cellular neighborhoods, bulk (cell type-agnostic) HLA expression along the tumor-stromal axis was initially plotted. This bulk expression was then further stratified by cell types, revealing patterns of ER dependence. Cell types were classified as ER-dependent or ER-independent based on their local expression profiles relative to ERα, calculated across pooled cells from all samples.

Additionally, a within-sample correlation analysis between ERα expression in cancer cells and HLA expression in all cell types was performed. This analysis complemented the pooled sample approach and highlighted distinct correlation patterns between ER-dependent and ER-independent cell types within the tumor microenvironment.

**ACOSOG clinical trial analysis**

The gene expression data from the ACOSOG Z1031B trial was subjected to quantile normalization and log-transformation to standardize expression levels across samples. Missing values (6% of the data) were imputed using a K-nearest neighbors (KNN) algorithm to ensure completeness before downstream analysis. Principal component analysis was applied to reduce the dimensionality of the expression data and investigate variability in relation to Ki-67 status changes (a marker of proliferation). Metadata including time points (pre- and post-treatment), Ki-67 status, and response groups (AI-resistant vs. sensitive) were utilized for the analysis, with response groups defined by further refined based on changes in Ki-67 from pre- to post-treatment.

Gene set enrichment was performed using ssGSEA to quantify the activity of key gene modules, such as the ERS and immune-related pathways, at both pre- and post-treatment stages. To capture broader pathway activity, FGSEA was performed using the ranked list of differentially expressed genes, combining both pre-annotated pathways hallmark pathways and custom modules. This approach highlighted significant differences in module activity between AI-sensitive and AI-resistant groups.

Two types of differential expression analyses were conducted: a baseline comparison between AI-resistant and sensitive groups, and a comparison of pre- to post-treatment changes for each group. Volcano plots were generated to visualize the DEGs, focusing on genes from the ERS and immune modules, as well as hallmark genes of interest (e.g., *ESR1, CD8A, HLA-A*). Finally, multivariate linear regression was used to identify factors contributing to changes in APM/TC module expression during treatment, providing insights into how ERS and immune pathways interact in the tumor microenvironment.

**Figure S1. Cross-cohort transcriptomic comparison in breast cancer (Related to Figure 1).**
**A.** Distribution of breast cancer subtypes across TCGA and METABRIC datasets.
**B.** Spearman correlation coefficients (SCC) between three hallmark genes' expression in HR+/HER2- samples from TCGA and METABRIC cohorts. *ESR1* was used as the hallmark gene for ERS, *HLA-A* for antigen presentation, and *CD8A* for T cells.
**C.** Comparison of mean expression of individual genes in TCGA and METABRIC (one point = one gene) with a fitted lowess curve (red). The horizontal dashed line indicates a signal threshold, expression level below which is harder to distinguish from noise in METABRIC microarray data.
**D**. Boxplot highlighting the distributions of mean absolute SCC across 778 genes in the curated gene list for TCGA and METABRIC cohorts. Significantly higher values in TCGA compared to METABRIC indicate that TCGA data is less affected by noise (paired two-tailed t-test).
**E.** Relationship between mean expression levels and mean absolute SCC among genes within each module in TCGA (left) and METABRIC (right). Genes with lower mean expression than the signal threshold (vertical line) in METABRIC showed particularly lower correlations due to noise, whereas TCGA data did not show the trend.
**F.** Relationship between cell cycle module and ERS module in HR+/HER2- samples from TCGA, assessed using gene set variation analysis (GSVA) z-scores for individual patients.
**G.** Distributions of SCC within ERS (left) and pan-immune modules (middle), and between ERS and pan-immune modules (right) in the TCGA cohort, grouped by breast cancer subtypes.
**H.** SCC within individual immune modules (TNFα/NF-κB, IFN-I, APM/TC) in the TCGA cohort.
**I.** SCC between genes in ERS and each immune module in the TCGA cohort. In panels **G-I**, orange and yellow indicate HR+ and HR- subtypes, respectively. To compute statistics, an equal number of patients were randomly selected from each subtype and a one sample two-tailed t-test was conducted for each population to assess the mean's deviation from zero (* $p < 1e-25$, ** $p < 1e-50$, *** $p < 1e-100$, **** $p < 1e-200$).
**J.** Heatmaps displaying SCC between 254 genes within ERS or three immune modules in four breast cancer subtypes.
**K.** Module-specific average SCC derived from **J**.

**Figure S2. HR+ dependency revealed in co-expression analysis across METABRIC (Related to Figure 1).**

**A.** Heatmap displaying SCC for 778 genes from the manually curated gene list in HR+/HER2- METABRIC patients, with indicators for hallmark genes (*ESR1*, *HLA-A*, *CD8A*), initial gene list (**Table S1**), and final gene modules (ERS, immune, or CC). Squares correspond to the three modules.

**B**. Heatmap showing SCC for 254 genes within the ERS or immune modules. Squares roughly correspond to individual immune panels.

**C.** Heatmaps showing SCC for 254 genes in four breast cancer subtypes from METABRIC data, emphasizing differential module interactions. Genes in the modules (ERS, TNFα/NF-κB, IFN-I, APM/TC) are shown in the same order to Figure **1B**.

**D**. Module-specific average SCC derived from **C**.

**E.** *ESR1* expression in four breast cancer subtypes from METABRIC data. The dashed line corresponds to the threshold distinguishing HR+ and HR- samples.

**F.** Relationship between *ESR1* gene expression and ERS module expression, with the *ESR1* threshold from **E**.

Two linear regression lines correspond to *ESR1*-low and *ESR1*-high populations, which largely correspond to HR- and HR+ subtypes.

**G**. Relationship between expression of ERS and three immune modules across all subtypes in METABRIC cohort. Two linear regression lines represent ERS-low and ERS-high populations, separated by vertical lines.

**H.** Distributions of SCC within ERS (left) and pan-immune modules (middle), and between ERS and pan-immune modules (right) in the METABRIC cohort, grouped by breast cancer subtypes.

**I.** SCC within individual immune modules (TNFα/NF-κB, IFN-I, APM/TC) in the METABRIC cohort.

**J.** SCC between genes in ERS and each immune module in the METABRIC cohort. In **H-J**, orange and yellow indicate HR+ and HR- subtypes. One sample two-tailed t-test was performed for each population to assess the mean's deviation from zero. To compute statistics, an equal number of patients were randomly selected from each subtype and a one sample two-tailed t-test was conducted for each population to assess the mean's deviation from zero (* $p < 1e-25$, ** $p < 1e-50$, *** $p < 1e-100$, **** $p < 1e-200$).

**K.** Relationship of TNFα/NF-κB and IFN-I module expression with APM/TC expression pooling all subtypes. Linear regression lines corresponding to ERS-low and ERS-high populations are shown.

**L.** Grouping of patients based on TNFα/NF-κB and IFN-I module expression (left) and corresponding APM/TC expression (right). Two linear regression lines in the left plot correspond to ERS-low and ERS-high populations. The samples were divided into four groups based on their TNFα/NF-κB and IFN-I expression compared with the median values.

**Figure S3. Validating module co-expressions in the MDACC cohort (Related to Figure 1).**
**A.** Heatmaps displaying SCC between genes that belong to either ERS or immune modules across 74 patients from the MD Anderson Cancer Center (MDACC) cohort. The samples were stratified into four breast cancer subtypes. Genes are shown in the same order to Figures **1B**.
**B.** Module-specific average correlation coefficients derived from **A**.

**Figure S4. Determining cell types in HR+ breast cancer on scRNA-seq data (Related to Figure 2).**
UMAP visualization of the single-cell RNA-seq data from 20 treatment-naïve HR+ primary breast cancer samples in WEHI cohort.
**A-B.** Overall cell population UMAP plots color-marked by patient identities (left), breast cancer subtypes (middle), and basality scores of cancer cells (right). Refer to the UMAP in Figure **2A** for the assigned cell types.
**B.** Dot plot highlighting lineage gene marker expression across identified cell types.
**C.** UMAP plots for the whole cell population, color-coded by the key cell type markers.
**D**. UMAP plots for lymphoid cells, color-coded by cell types and lineage biomarker expression.
**E**. UMAP plots for myeloid cells, color-coded by cell types and lineage biomarker expression.

**Figure S5. Module expression analysis of scRNA-seq data in the WEHI cohort (Related to Figure 2).**

**A**. Relationship between *ESR1* gene expression and ERS module expression in cancer cells in the WEHI cohort. The expression was averaged per patient.

**B**. Comparative analysis of module expression profiles for each cell type.

**C.** PCC between patient-level module expression profiles across HR+ subtypes (left) and all subtypes (right).

**D**. Co-occurrence between cell type frequency among 34 patients representing all subtypes in the WEHI cohort.

**E.** Correlation of ERS module expression per patient with each cell type's frequency among 34 patients representing all subtypes.

**Figure S6. Identification of conserved module expression in breast cancer cell lines (Related to Figure 3).**
**A**. Scatterplot comparing the expression of module genes in cancer cells from patient samples (WEHI) versus CCLE cell lines. Each point represents a single gene. The horizontal line indicates the signal threshold; genes below this line are considered not expressed in cell lines.
**B.** Expression levels of genes in cancer cell lines, grouped by the cell types that most dominantly expressed them in the WEHI dataset (Figure **2E,** middle panel). Each point corresponds to a single gene. The horizontal line marks the signal threshold, distinguishing genes expressed in cell lines from those not expressed.
**C**. Pairwise SCC of gene expression across 54 breast cancer cell lines, covering all PAM50 subtypes. Three boxes highlight three cancer cell-intrinsic modules.
**D**. Module-specific average SCC values among 54 breast cancer cell lines, derived from **C**.
**E.** List of genes in cancer cell-intrinsic gene modules. Immune modules are further sub-divided into "sub" modules based on the original modules derived from the whole-tumor analysis.
**F.** Pathway enrichment analysis against intrinsic TNFα/NF-κB and IFN-I modules. (i) First, the enrichment of 8,077 gene sets that belong to Hallmark, GO, KEGG, Reactome, and Biocarta, were assessed against correlated and non-correlated genes. 26 pathways enriched in 'non-correlated' genes (top-left plot; FDR-adjusted p-values < 1e-3) were excluded. (ii) The remaining pathways were analyzed for enrichment in intrinsic TNFα/NF-κB and IFN-I modules, identifying 1 and 21 pathways enriched for TNFα/NF-κB and IFN-I modules, respectively (top-right plot; FDR-adjusted p-values < 1e-5). (iii) Of the 21 pathways enriched in the intrinsic IFN-I module, 13 and 4 were enriched in APM_sub and IFN_sub modules, (bottom-right; FDR-adjusted p-values < 1e-5).
**G.** The table of enriched pathways and their corresponding -log10(p-values) from the analysis in panel **F**. Enrichment was assessed using a one-sided Fisher's exact test. Ag: antigen, AgPP: Ag processing and presentation.
**H**. Expression for individual genes in the cancer cell-intrinsic ERS, TNFα/NF-κB, and IFN-I modules across 54 cell lines. PAM50 classification for each cell line is also shown at the top.

**Figure S7. Module expression analysis of snRNA-seq data in the DF/BCC cohort (Related to Figure 4).**
**A.** Heatmaps showing the SCC for individual genes in the ERS and immune modules across 29 samples.
**B**. Expression of the four gene modules across 29 samples, computed with single-sample GSEA on bulk RNA-seq data.
**C.** PCC among expression of the four modules across 29 samples.
**D.** UMAP representation of the whole cell population from 17 samples in the DF/BCC cohort.
**E.** Dot plot highlighting lineage marker expression in the identified cell types.
**F.** Relationship between mean *ESR1* gene expression in cancer cells from snRNA-seq data and ERS module expression in cancer cells from bulk RNA-seq data per patient in the WEHI cohort.
**G.** Expression of the four modules at the single-cell level, pooling data from 29 samples.
**H.** Heatmap of mean module expression (ssGSEA enrichment scores) by cell type and by sample.
**I.** Deconvolution of module expression from snRNA-seq data to patient-level and cell type-level expression.
**J.** PCC between the per-patient level expression profiles of the ERS and immune modules.
**K.** Cell type frequency of 17 samples reconstituted from the snRNA-seq data. Samples are sorted by ERS module expression calculated using bulk RNA-seq data.
**L**. Cell type co-occurrence computed by PCC across 17 samples in the snRNA-seq data.
**M**. Bar plot showing the Pearson correlation coefficients of cell type frequencies with ERS module expression across 17 samples.

**Figure S8. Cell type calling in CyCIF and correlation of cancer cell-intrinsic IFN-I module expression with cancer cell frequency (Related to Figure 4).**
Schematic cell type definition tree displaying lineage markers used to identify corresponding cell types and the list of cell state markers analyzed in CyCIF.

**Figure S9. Tumor-to-stroma ratio analysis in CyCIF reveals the relationship between spatial distribution of cell types and module expression (Related to Figure 5).**
**A.** Micrograph illustrating the colocalization of CD4+ and CD8+ T cells with CD163+ macrophages (Mac_CD163 and Mac_CD68_CD163) in patient ID df27 (ERS-low tumor). The bottom row shows a magnified view of the area highlighted by the square in the top row.
**B.** Mean PD-1 expression per T cell subset along tumor-stroma axis.
**C.** Relationship between T cell density (# of T cells/$mm^2$) and mean PD-1 expression across 28 samples.
**D.** Relationship between the TS ratio of Mac_CD163 and TNFα/NF-κB (left), IFN-I (middle), and APM/TC (right) module expression across 28 samples. Positive correlations indicate that higher module expression is associated with greater stromal prevalence of Mac_CD163.
**E.** Tumor-to-stroma (TS) ratio of three macrophage subsets. TS ratios of different subsets from each sample are connected by lines. Paired two-sided t-test was performed to assess significance.
**F.** Relationship between the TS ratio of endothelial cells and TNFα/NF-κB expression. Positive correlations indicate that higher module expression is associated with greater stromal prevalence of endothelial cells.

**Figure S10. Spatial analysis of HLA expression reveals ER-dependent and ER-independent HLA expression patterns (Related to Figure 6).**

**A.** Comparison of protein expression between the tumor core and stroma for ERα, HLA-ABC, and HLA-DPB1. Protein expression was averaged across regions 150-300μm from the tumor border in both tumor core and stroma. Multivariate linear regression was applied to evaluate (1) the relationship between T cell density and individual protein expression, and (2) differences in protein expression between the tumor core and stroma.

**B.** Pearson correlation coefficients between mean expression per cell type with ERS transcriptional activity across samples.

**C.** Average expression of HLA-ABC (top-left) and HLA-DPB1 (top-right) in each sample, grouped by cell types. Cell types are further stratified by the average expression across samples. A paired two-tailed t-test was performed for all combinations of cell types (bottom-left).

**D.** CN clusters sorted by their abundance in ERS-high samples versus ERS-low samples. Top: Barplot of SCC between CN cluster frequency and ERS module expression across 28 samples. Bottom: heatmap of relative frequency of 100 CN clusters within each sample.

**E.** Heatmap highlighting a hierarchical clustering of mean ERα, HLA-ABC, and HLA-DPB1 expression patterns per cell type across samples. The patterns were grouped by proteins primarily rather than cell types.

**Figure S11 Supporting data for module expression analysis on ACOSOG Z1031B trial data (Related to Figure 7).**

**A.** Percentage of Ki67-positive tumors before and after treatment. Ki67 levels increased in 34 resistant patients (blue) but significantly decreased in the 34 most sensitive patients (red), selected based on the greatest reduction in Ki67 levels (post-treatment divided by pre-treatment).
**B.** Relationship between module expression at pre-treatment and changes in module expressions during treatment. For each module (diagonal), Spearman correlation coefficients were negative, indicating that samples with higher ERS module expression or lower immune module expression at baseline showed bigger decrease in ΔERS or increase in ΔTNFα/NF-κB, ΔIFN-I, and ΔAPM/TC, respectively. This aligns with the Figure **7A,B**.
**C**. SCC between module expression and cell type abundance at pre-treatment.
**D**. Relationship between the TNFα/NF-κB module expression and the endothelial cell abundance at pre-treatment.
**E**. Relationship between the changes in TNFα/NF-κB module expression and endothelial cells during AI treatment.

**Table S1. Curated gene list with references.** This table provides a list of genes associated with estrogen receptor signaling (ERS), antigen presentation (APM), and T cell (TC) activation, including the references from which these genes were compiled.

**Table S2. Gene in each module.** This table details the five gene modules identified through analysis of bulk transcriptome data from TCGA and METABRIC cohorts, namely ERS, CC (cell cycle), TNFα/NF-κB, IFN-I, and APM/TC modules. This table is related to **Fig. 1B,C.**

**Table S3. Clinical metadata for samples from the DF/BCC cohort.** This table provides the clinical metadata associated with 29 samples analyzed in this study.

**Supplementary Data S1. Interactive view of CyCIF staining of an ERS-high tumor (Patient: df2 in DF/BCC cohort)**

**https://www.cycif.org/data/shimada-2024/ERS_high**

**Supplementary Data S2. Interactive view of CyCIF staining of an ERS-low tumor (Patient: df27 in DF/BCC cohort)**

**https://www.cycif.org/data/shimada-2024/ERS_low**
