## Supplementary Figures for "An estrogen receptor signaling transcriptional program linked to immune evasion in human hormone receptor-positive breast cancer"

### Figure S1

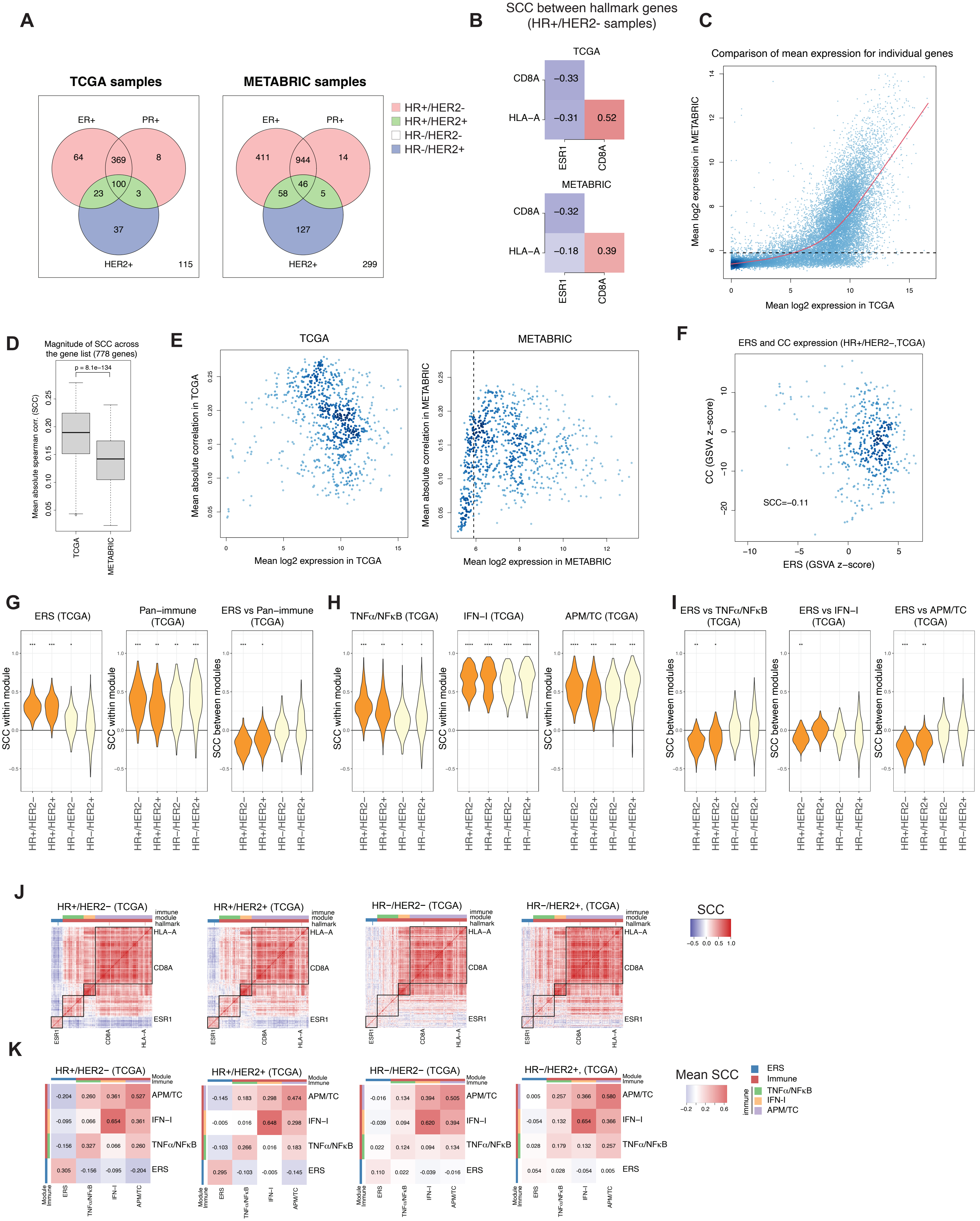

A
TCGA samples
ER+
PR+
64
369
8
100
23
3
37
HER2+
115
METABRIC samples
ER+
PR+
411
944
14
46
58
5
127
HER2+
299
HR+/HER2-
HR+/HER2+
HR-/HER2-
HR-/HER2+
B
SCC between hallmark genes
(HR+/HER2- samples)
TCGA
CD8A
−0.33
HLA−A
−0.31
0.52
ESR1
CD8A
METABRIC
CD8A
−0.32
HLA−A
−0.18
0.39
ESR1
CD8A
C
Comparison of mean expression for individual genes
Mean log2 expression in METABRIC
Mean log2 expression in TCGA
D
Magnitude of SCC across
the gene list (778 genes)
p = 8.1e−134
Mean absolute spearman corr. (SCC)
TCGA
METABRIC
E
TCGA
Mean absolute correlation in TCGA
Mean log2 expression in TCGA
METABRIC
Mean absolute correlation in METABRIC
Mean log2 expression in METABRIC
F
ERS and CC expression (HR+/HER2−,TCGA)
CC (GSVA z−score)
SCC=−0.11
ERS (GSVA z−score)
G
ERS (TCGA)
Pan−immune
(TCGA)
ERS vs Pan−immune
(TCGA)
SCC within module
SCC between modules
HR+/HER2−
HR+/HER2+
HR−/HER2−
HR−/HER2+
H
TNFα/NFκB (TCGA)
IFN−I (TCGA)
APM/TC (TCGA)
I
ERS vs TNFα/NFκB
(TCGA)
ERS vs IFN−I
(TCGA)
ERS vs APM/TC
(TCGA)
J
HR+/HER2− (TCGA)
HR+/HER2+ (TCGA)
HR−/HER2− (TCGA)
HR−/HER2+, (TCGA)
immune
module
hallmark
HLA−A
CD8A
ESR1
SCC
K
HR+/HER2− (TCGA)
-0.204 0.260 0.361 0.527 APM/TC
-0.095 0.066 0.654 0.361 IFN−I
-0.156 0.327 0.066 0.260 TNFα/NFκB
0.305 -0.156 -0.095 -0.204 ERS
HR+/HER2+ (TCGA)
-0.145 0.183 0.298 0.474 APM/TC
-0.005 0.016 0.648 0.298 IFN−I
-0.103 0.266 0.016 0.183 TNFα/NFκB
0.295 -0.103 -0.005 -0.145 ERS
HR−/HER2− (TCGA)
-0.016 0.134 0.394 0.505 APM/TC
-0.039 0.094 0.620 0.394 IFN−I
0.022 0.124 0.094 0.134 TNFα/NFκB
0.110 0.022 -0.039 -0.016 ERS
HR−/HER2+, (TCGA)
0.005 0.257 0.366 0.580 APM/TC
-0.054 0.132 0.654 0.366 IFN−I
0.028 0.179 0.132 0.257 TNFα/NFκB
0.054 0.028 -0.054 0.005 ERS
Module
Immune
ERS
TNFα/NFκB
IFN−I
APM/TC
Mean SCC
ERS
Immune
TNFα/NFκB
IFN-I
APM/TC
immune

### Figure S2

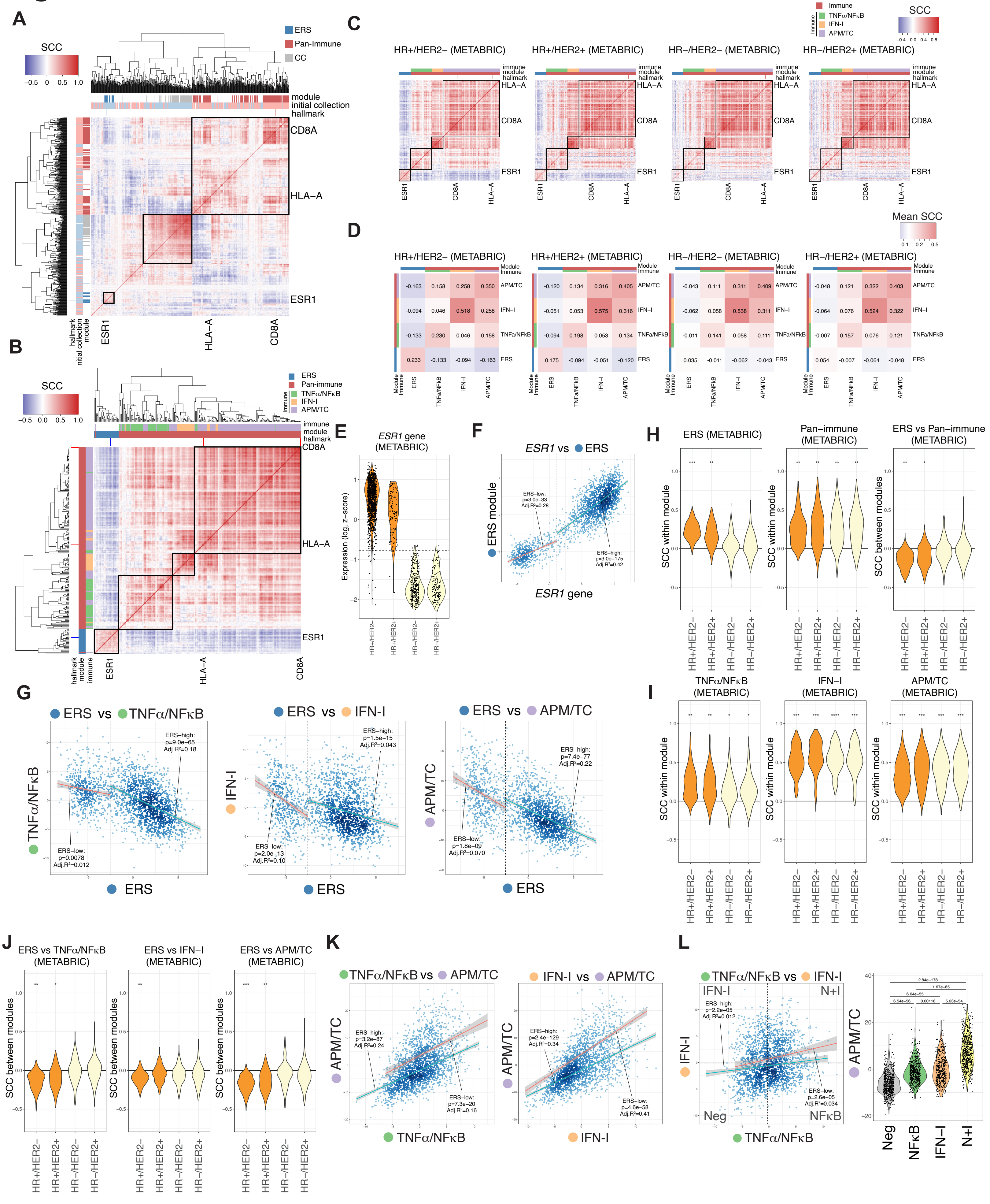
A
SCC
−0.5 0 0.5 1.0
ERS
Pan-Immune
CC
module
initial collection
hallmark
CD8A
HLA−A
ESR1
B
SCC
−0.5 0.0 0.5 1.0
ERS
Pan-immune
Immune
TNFα/NFκB
IFN-I
APM/TC
immune
module
hallmark
CD8A
HLA−A
ESR1
C
HR+/HER2− (METABRIC)
HR+/HER2+ (METABRIC)
HR−/HER2− (METABRIC)
HR−/HER2+ (METABRIC)
Immune
D
Mean SCC
HR+/HER2− (METABRIC)
-0.163 0.158 0.258 0.350 APM/TC
-0.094 0.046 0.518 0.258 IFN−I
-0.133 0.230 0.046 0.158 TNFa/NFκB
0.233 -0.133 -0.094 -0.163 ERS
HR+/HER2+ (METABRIC)
-0.120 0.134 0.316 0.405 APM/TC
-0.051 0.053 0.575 0.316 IFN−I
-0.094 0.198 0.053 0.134 TNFa/NFκB
0.175 -0.094 -0.051 -0.120 ERS
HR−/HER2− (METABRIC)
-0.043 0.111 0.311 0.409 APM/TC
-0.062 0.058 0.538 0.311 IFN−I
-0.011 0.141 0.058 0.111 TNFa/NFκB
0.035 -0.011 -0.062 -0.043 ERS
HR−/HER2+ (METABRIC)
-0.048 0.121 0.322 0.403 APM/TC
-0.064 0.076 0.524 0.322 IFN−I
-0.007 0.157 0.076 0.121 TNFa/NFκB
0.054 -0.007 -0.064 -0.048 ERS
Module
Immune
E
ESR1 gene
(METABRIC)
Expression (log2 z−score)
HR+/HER2−
HR+/HER2+
HR−/HER2−
HR−/HER2+
F
ESR1 vs ERS
ERS module
ESR1 gene
ERS−low:
p=3.0e−33
Adj.R²=0.28
ERS−high:
p=3.0e−175
Adj.R²=0.42
G
ERS vs TNFα/NFκB
ERS−high:
p=9.0e−65
Adj.R²=0.18
ERS−low:
p=0.0078
Adj.R²=0.012
ERS vs IFN−I
ERS−high:
p=1.5e−15
Adj.R²=0.043
ERS−low:
p=2.0e−13
Adj.R²=0.10
ERS vs APM/TC
ERS−high:
p=7.4e−77
Adj.R²=0.22
ERS−low:
p=1.8e−09
Adj.R²=0.070
H
ERS (METABRIC)
Pan−immune
(METABRIC)
ERS vs Pan−immune
(METABRIC)
SCC within module
SCC between modules
I
TNFα/NFκB
(METABRIC)
IFN−I
(METABRIC)
APM/TC
(METABRIC)
SCC within module
J
ERS vs TNFα/NFκB
(METABRIC)
ERS vs IFN−I
(METABRIC)
ERS vs APM/TC
(METABRIC)
SCC between modules
K
TNFα/NFκB vs APM/TC
ERS−high:
p=3.2e−87
Adj.R²=0.24
ERS−low:
p=7.3e−20
Adj.R²=0.16
IFN-I vs APM/TC
ERS−high:
p=2.4e−129
Adj.R²=0.34
ERS−low:
p=4.6e−58
Adj.R²=0.41
L
TNFα/NFκB vs IFN-I
IFN-I
N+I
Neg
NFκB
ERS−high:
p=2.2e−05
Adj.R²=0.012
ERS−low:
p=2.6e−05
Adj.R²=0.034
APM/TC
2.84e−178
1.67e−85
6.64e−55
6.54e−56
0.00118
5.63e−54
Neg
NFκB
IFN−I
N+I

### Figure S3

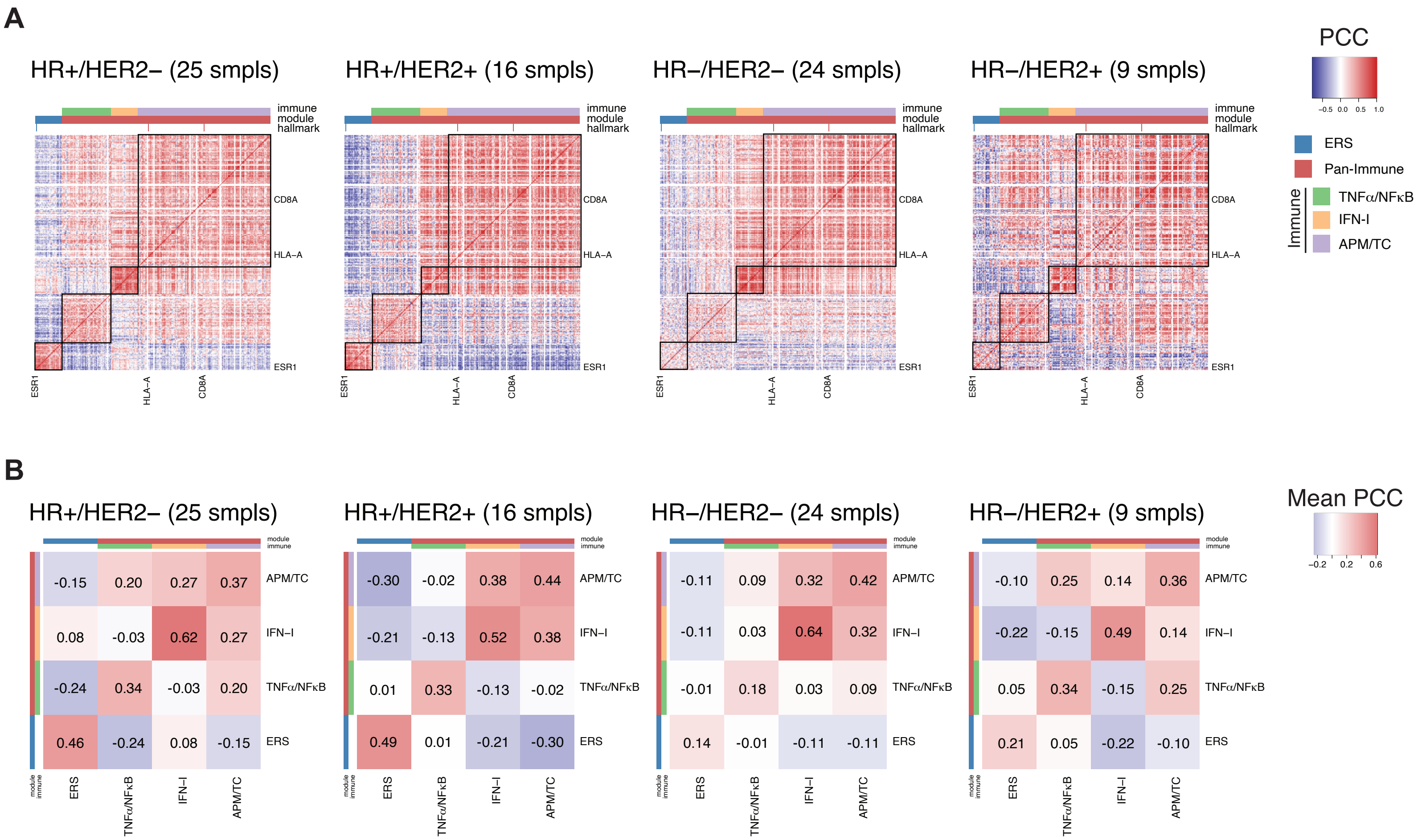
A
HR+/HER2− (25 smpls)
HR+/HER2+ (16 smpls)
HR−/HER2− (24 smpls)
HR−/HER2+ (9 smpls)
immune
module
hallmark
CD8A
HLA−A
ESR1
PCC
−0.5 0.0 0.5 1.0
ERS
Pan-Immune
Immune
TNFα/NFκB
IFN-I
APM/TC
B
Mean PCC
−0.2 0.2 0.6
module
immune
APM/TC
IFN−I
TNFα/NFκB
ERS
HR+/HER2− (25 smpls): APM/TC -0.15 0.20 0.27 0.37; IFN−I 0.08 -0.03 0.62 0.27; TNFα/NFκB -0.24 0.34 -0.03 0.20; ERS 0.46 -0.24 0.08 -0.15
HR+/HER2+ (16 smpls): APM/TC -0.30 -0.02 0.38 0.44; IFN−I -0.21 -0.13 0.52 0.38; TNFα/NFκB 0.01 0.33 -0.13 -0.02; ERS 0.49 0.01 -0.21 -0.30
HR−/HER2− (24 smpls): APM/TC -0.11 0.09 0.32 0.42; IFN−I -0.11 0.03 0.64 0.32; TNFα/NFκB -0.01 0.18 0.03 0.09; ERS 0.14 -0.01 -0.11 -0.11
HR−/HER2+ (9 smpls): APM/TC -0.10 0.25 0.14 0.36; IFN−I -0.22 -0.15 0.49 0.14; TNFα/NFκB 0.05 0.34 -0.15 0.25; ERS 0.21 0.05 -0.22 -0.10

### Figure S4

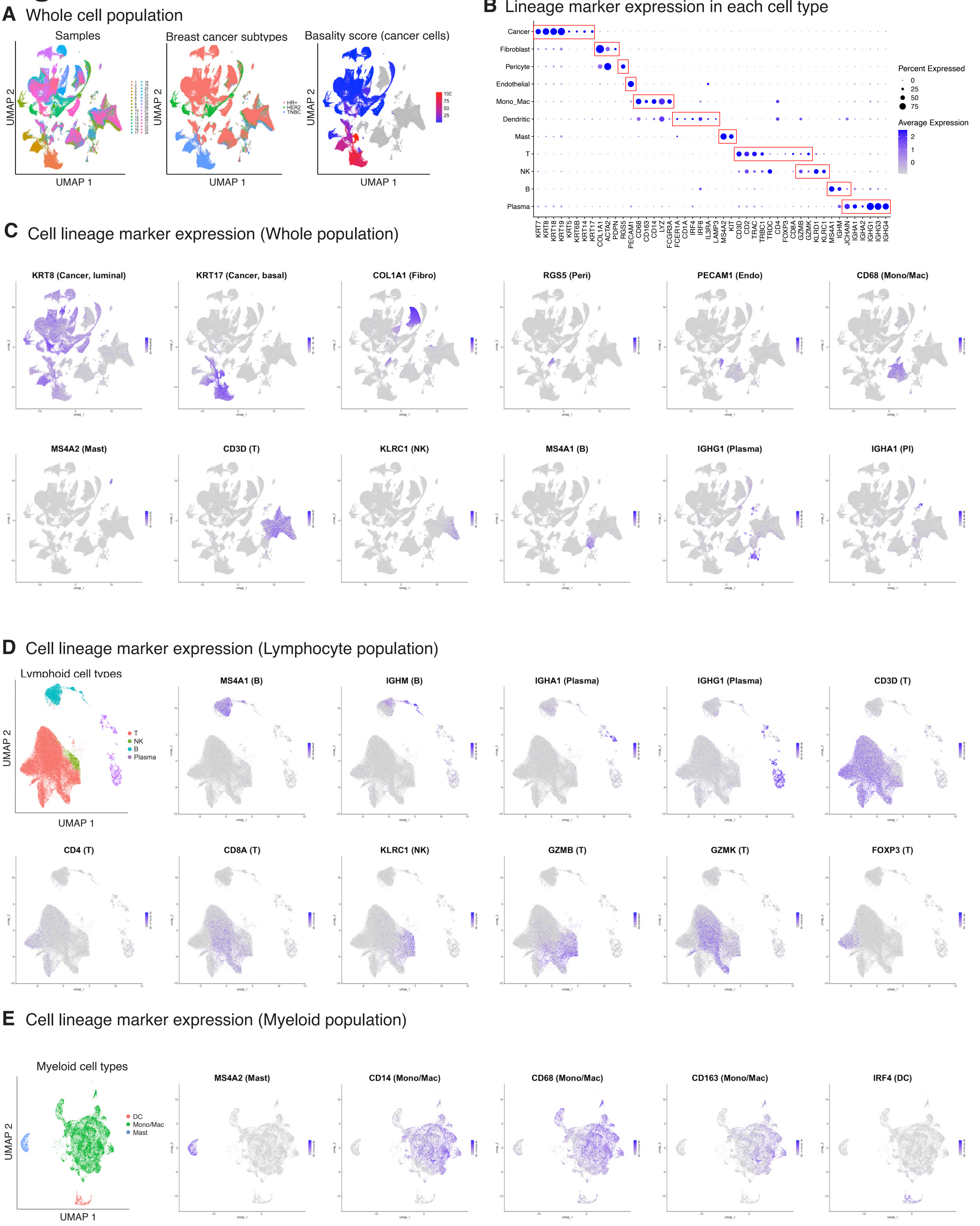

A Whole cell population
Samples
Breast cancer subtypes
Basality score (cancer cells)
UMAP 1
UMAP 2
HR+
HER2
TNBC
B Lineage marker expression in each cell type
Cancer
Fibroblast
Pericyte
Endothelial
Mono_Mac
Dendritic
Mast
T
NK
B
Plasma
KRT7
KRT8
KRT18
KRT19
KRT5
KRT6B
KRT14
KRT17
COL1A1
ACTA2
PDPN
RGS5
PECAM1
CD68
CD163
CD14
LYZ
FCGR3A
FCER1A
CD1A
IRF4
IRF8
IL3RA
LAMP3
MS4A2
KIT
CD3D
CD2
TRAC
TRBC1
TRDC
CD4
FOXP3
CD8A
GZMB
GZMK
KLRD1
KLRC1
MS4A1
IGHM
JCHAIN
IGHA1
IGHA2
IGHG1
IGHG3
IGHG4
Percent Expressed
0
25
50
75
Average Expression
2
1
0
C Cell lineage marker expression (Whole population)
KRT8 (Cancer, luminal)
KRT17 (Cancer, basal)
COL1A1 (Fibro)
RGS5 (Peri)
PECAM1 (Endo)
CD68 (Mono/Mac)
MS4A2 (Mast)
CD3D (T)
KLRC1 (NK)
MS4A1 (B)
IGHG1 (Plasma)
IGHA1 (Pl)
D Cell lineage marker expression (Lymphocyte population)
Lymphoid cell types
T
NK
B
Plasma
UMAP 1
UMAP 2
MS4A1 (B)
IGHM (B)
IGHA1 (Plasma)
IGHG1 (Plasma)
CD3D (T)
CD4 (T)
CD8A (T)
KLRC1 (NK)
GZMB (T)
GZMK (T)
FOXP3 (T)
E Cell lineage marker expression (Myeloid population)
Myeloid cell types
DC
Mono/Mac
Mast
UMAP 1
UMAP 2
MS4A2 (Mast)
CD14 (Mono/Mac)
CD68 (Mono/Mac)
CD163 (Mono/Mac)
IRF4 (DC)

### Figure S5

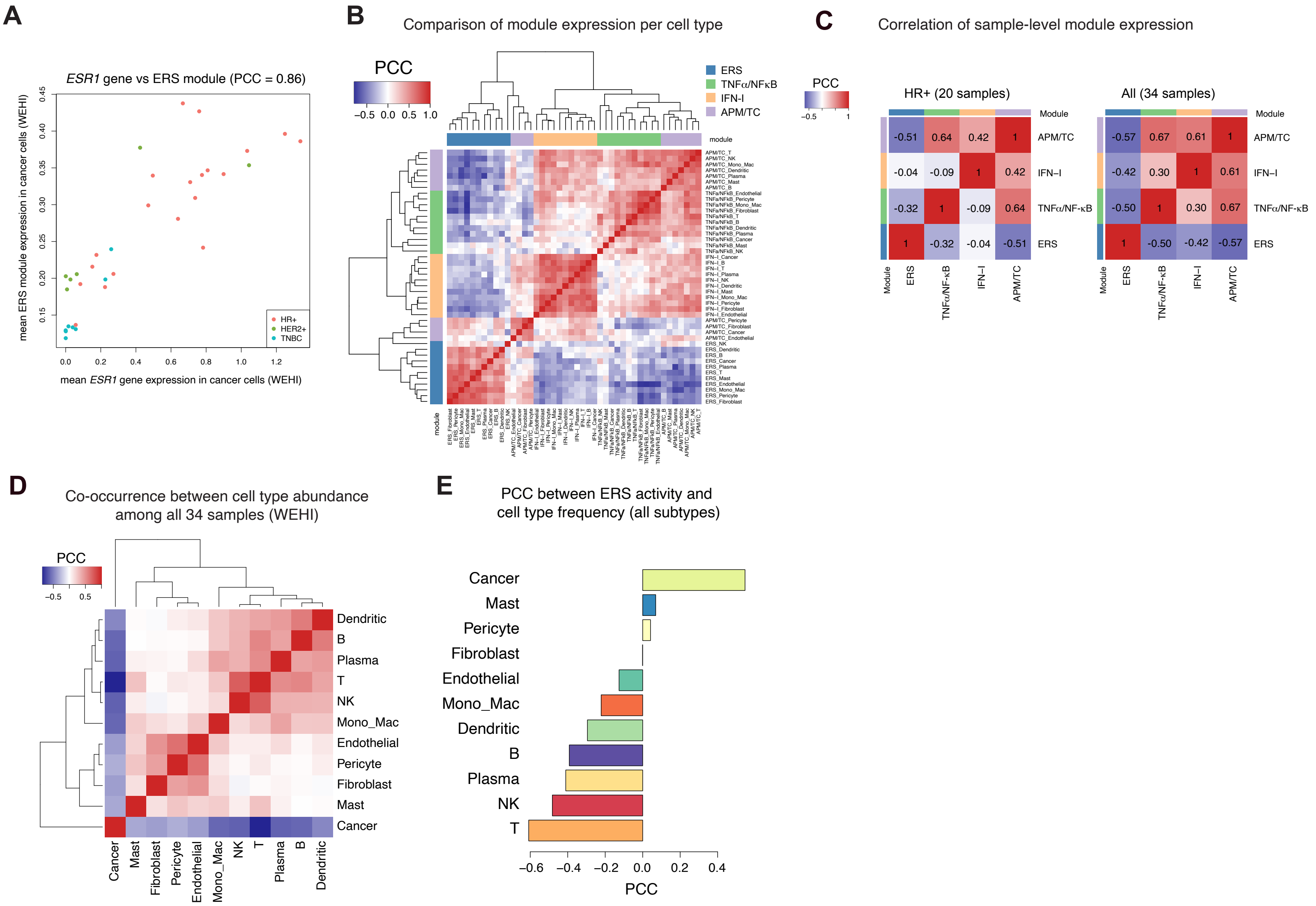
A
ESR1 gene vs ERS module (PCC = 0.86)
mean ERS module expression in cancer cells (WEHI)
mean ESR1 gene expression in cancer cells (WEHI)
HR+
HER2+
TNBC
B
Comparison of module expression per cell type
PCC
ERS
TNFα/NFκB
IFN-I
APM/TC
module
C
Correlation of sample-level module expression
PCC
HR+ (20 samples)
All (34 samples)
Module
APM/TC
IFN−I
TNFα/NF-κB
ERS
D
Co-occurrence between cell type abundance among all 34 samples (WEHI)
PCC
Dendritic
B
Plasma
T
NK
Mono_Mac
Endothelial
Pericyte
Fibroblast
Mast
Cancer
E
PCC between ERS activity and cell type frequency (all subtypes)
Cancer
Mast
Pericyte
Fibroblast
Endothelial
Mono_Mac
Dendritic
B
Plasma
NK
T
PCC

### Figure S6

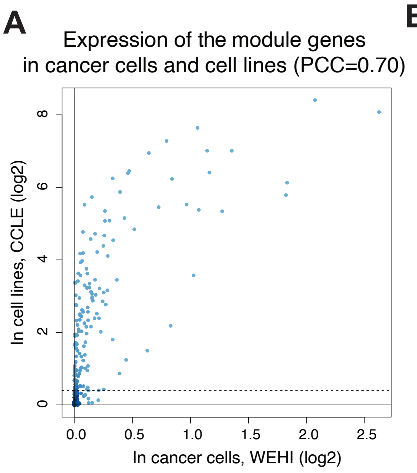


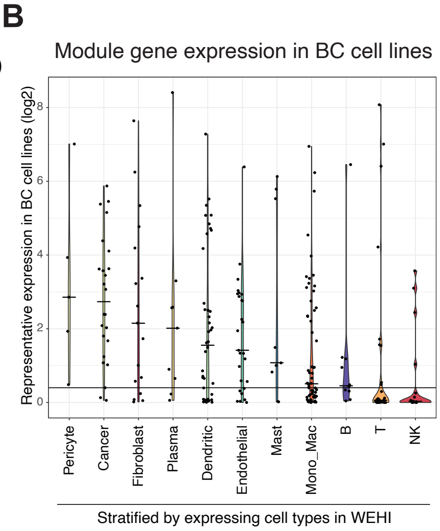


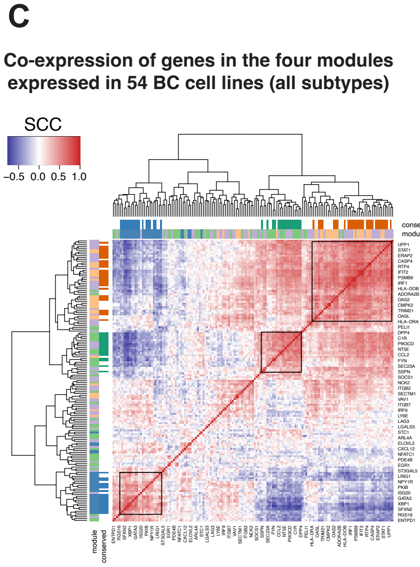


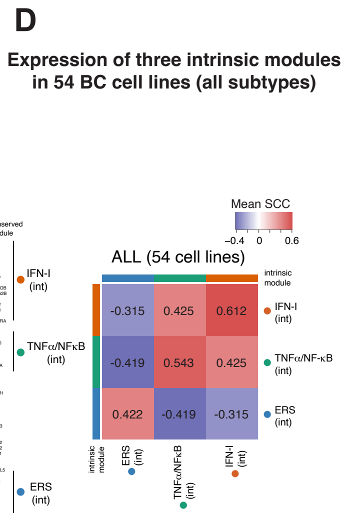


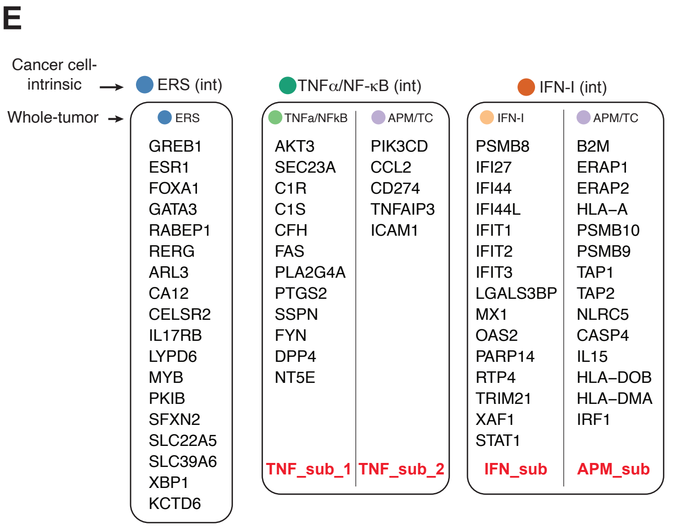


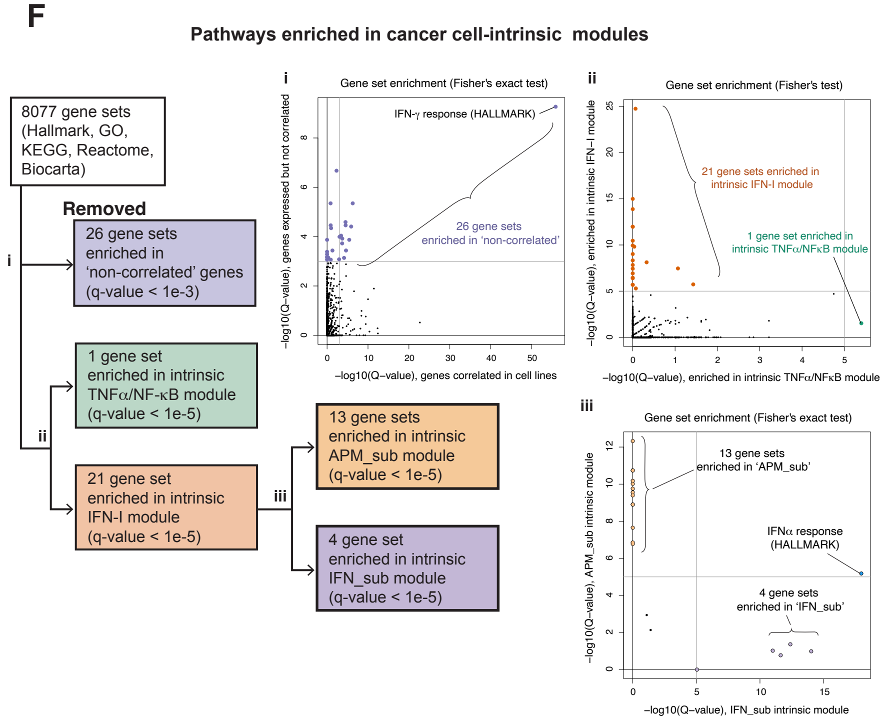


G

| Gene sets | TNFα/NFκB (int) | TNF_sub_1 (within TNFα/NFκB (int)) | TNF_sub_2 (within TNFα/NFκB (int)) | IFN-I | IFN_sub (within IFN_sub (int)) | APM_sub (within IFN_sub (int)) |
|---|---|---|---|---|---|---|
| Complement (Hallmark) | 5.40 | 4.80 | 0.00 | 1.52 | 0.00 | 2.48 |
| IFNα response (Hallmark) | 0.07 | 0.03 | 0.00 | 24.76 | 17.97 | 5.18 |
| IFNα/β signaling (Reactome) | 0.00 | 0.00 | 0.00 | 14.99 | 14.03 | 0.99 |
| IFN signaling (Reactome) | 0.00 | 0.00 | 0.00 | 13.88 | 12.39 | 1.36 |
| Response to virus (GOBP) | 0.00 | 0.00 | 0.00 | 11.97 | 11.63 | 0.77 |
| Defense response to symbiont (GOBP) | 0.00 | 0.00 | 0.00 | 11.97 | 11.00 | 1.02 |
| Response to type I IFN (GOBP) | 0.00 | 0.00 | 0.00 | 5.68 | 5.05 | 0.00 |
| AgPP of peptide antigen (GOBP) | 0.00 | 0.00 | 0.00 | 9.94 | 0.00 | 12.32 |
| AgPP (GOBP) | 0.04 | 0.00 | 0.11 | 9.81 | 0.00 | 10.74 |
| AgPP of endogenous peptide Ag (GOBP) | 0.00 | 0.00 | 0.00 | 9.02 | 0.00 | 10.74 |
| AgPP of endogenous Ag (GOBP) | 0.00 | 0.00 | 0.00 | 8.33 | 0.00 | 10.18 |
| Ag pres., folding assembly, pep loading of MHC-I (Reactome) | 0.33 | 0.23 | 0.00 | 8.12 | 0.00 | 10.01 |
| AgPP of pep Ag via MHC class I (GOBP) | 0.00 | 0.00 | 0.00 | 7.83 | 0.00 | 9.75 |
| Allograft rejection (Hallmark) | 1.07 | 0.00 | 0.66 | 7.46 | 0.00 | 8.90 |
| AgPP of exogenous peptide Ag (GOBP) | 0.00 | 0.00 | 0.00 | 7.43 | 0.00 | 9.40 |
| AgPP of exogenous Ag (GOBP) | 0.00 | 0.00 | 0.00 | 6.92 | 0.00 | 8.90 |
| Ag processing-cross presentation (Reactome) | 0.00 | 0.00 | 0.00 | 6.45 | 0.00 | 6.85 |
| Class I MHC mediated AgPP (Reactome) | 0.00 | 0.00 | 0.00 | 6.40 | 0.00 | 6.77 |
| AgPP (KEGG) | 0.00 | 0.00 | 0.00 | 5.68 | 0.00 | 7.65 |
| MHC pathway (Biocarta) | 0.00 | 0.00 | 0.00 | 10.46 | 0.00 | 9.55 |
| Regulation of response to biotic stimulus (GOBP) | 1.43 | 0.23 | 0.46 | 5.73 | 1.10 | 2.94 |
| IFNγ signaling (Reactome) | 0.07 | 0.00 | 0.13 | 5.29 | 1.41 | 2.13 |

Numerical values are -log P values with one-sided Fisher's exact test. Values larger than 5 are shown in red.

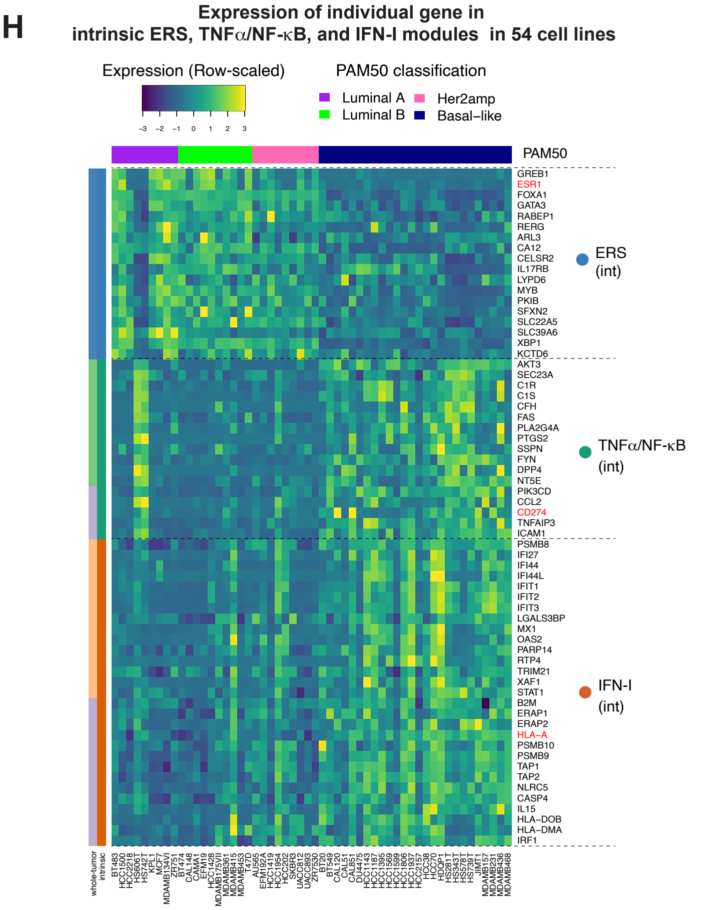

### Figure S7

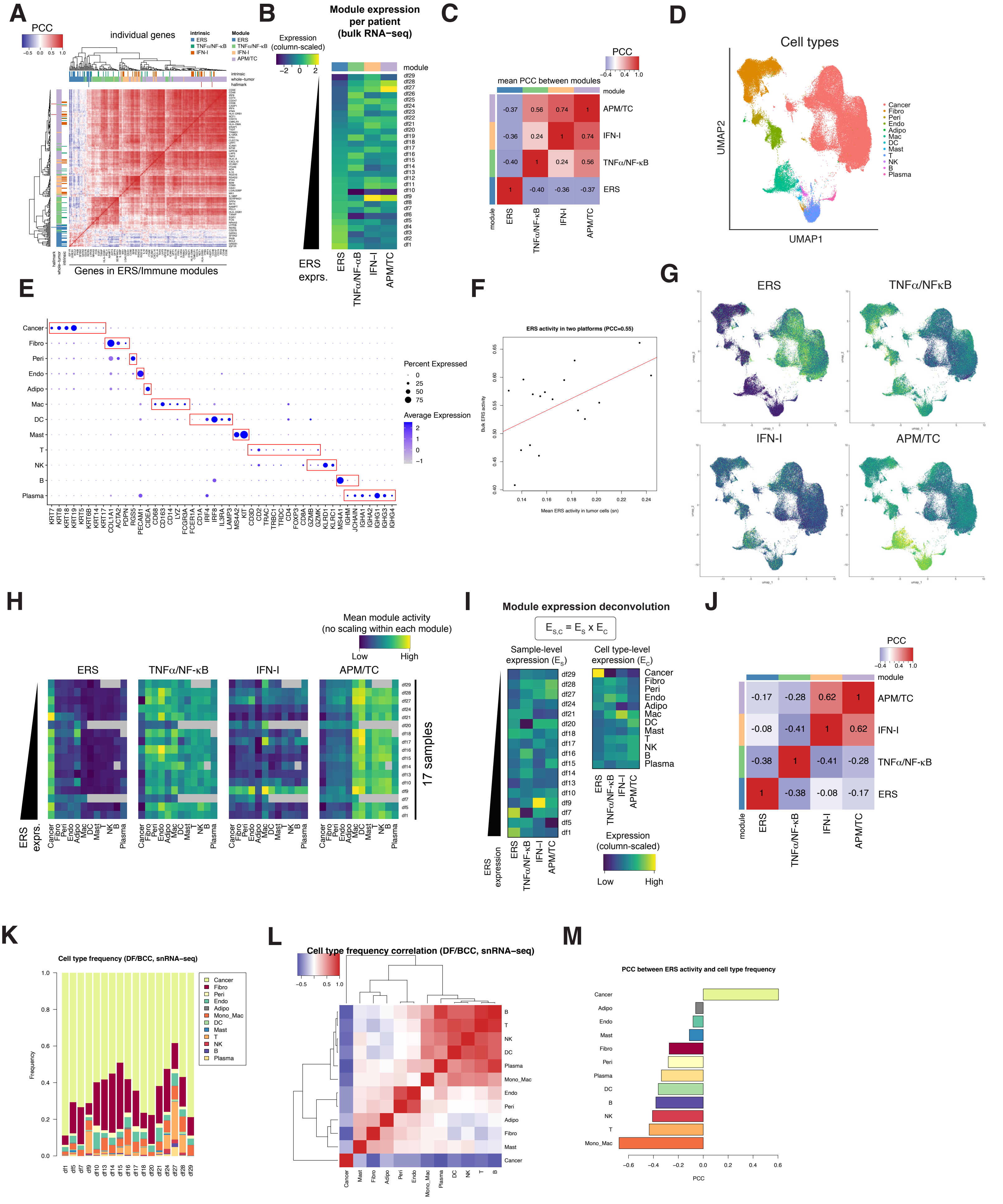

A
PCC
individual genes
intrinsic
ERS
TNFα/NF-κB
IFN-I
Module
ERS
TNFα/NF-κB
IFN-I
APM/TC
intrinsic
whole-tumor
hallmark
Genes in ERS/Immune modules
B
Module expression per patient (bulk RNA-seq)
Expression (column-scaled)
module
ERS exprs.
ERS
TNFα/NF-κB
IFN-I
APM/TC
C
PCC
mean PCC between modules
module
APM/TC
IFN-I
TNFα/NF-κB
ERS
D
Cell types
UMAP2
UMAP1
Cancer
Fibro
Peri
Endo
Adipo
Mac
DC
Mast
T
NK
B
Plasma
E
Percent Expressed
Average Expression
F
ERS activity in two platforms (PCC=0.55)
Bulk ERS activity
Mean ERS activity in tumor cells (sn)
G
ERS
TNFα/NFκB
IFN-I
APM/TC
H
Mean module activity (no scaling within each module)
Low
High
ERS exprs.
17 samples
I
Module expression deconvolution
Sample-level expression ($E_S$)
Cell type-level expression ($E_C$)
ERS expression
Expression (column-scaled)
Low
High
J
PCC
module
K
Cell type frequency (DF/BCC, snRNA−seq)
Frequency
Mono_Mac
L
Cell type frequency correlation (DF/BCC, snRNA−seq)
M
PCC between ERS activity and cell type frequency
PCC

### Figure S8

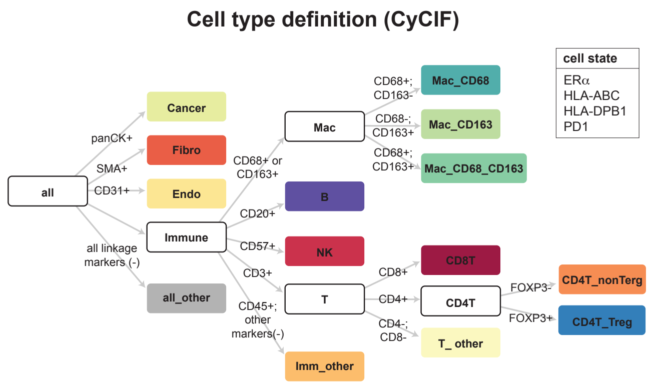
Cell type definition (CyCIF)
cell state
ERα
HLA-ABC
HLA-DPB1
PD1
all
panCK+
SMA+
CD31+
all linkage markers (-)
Cancer
Fibro
Endo
Immune
all_other
CD68+ or CD163+
CD20+
CD57+
CD3+
CD45+; other markers(-)
Mac
B
NK
T
Imm_other
CD68+; CD163-
CD68-; CD163+
CD68+; CD163+
Mac_CD68
Mac_CD163
Mac_CD68_CD163
CD8+
CD4+
CD4-; CD8-
CD8T
CD4T
T_ other
FOXP3-
FOXP3+
CD4T_nonTerg
CD4T_Treg

### Figure S9

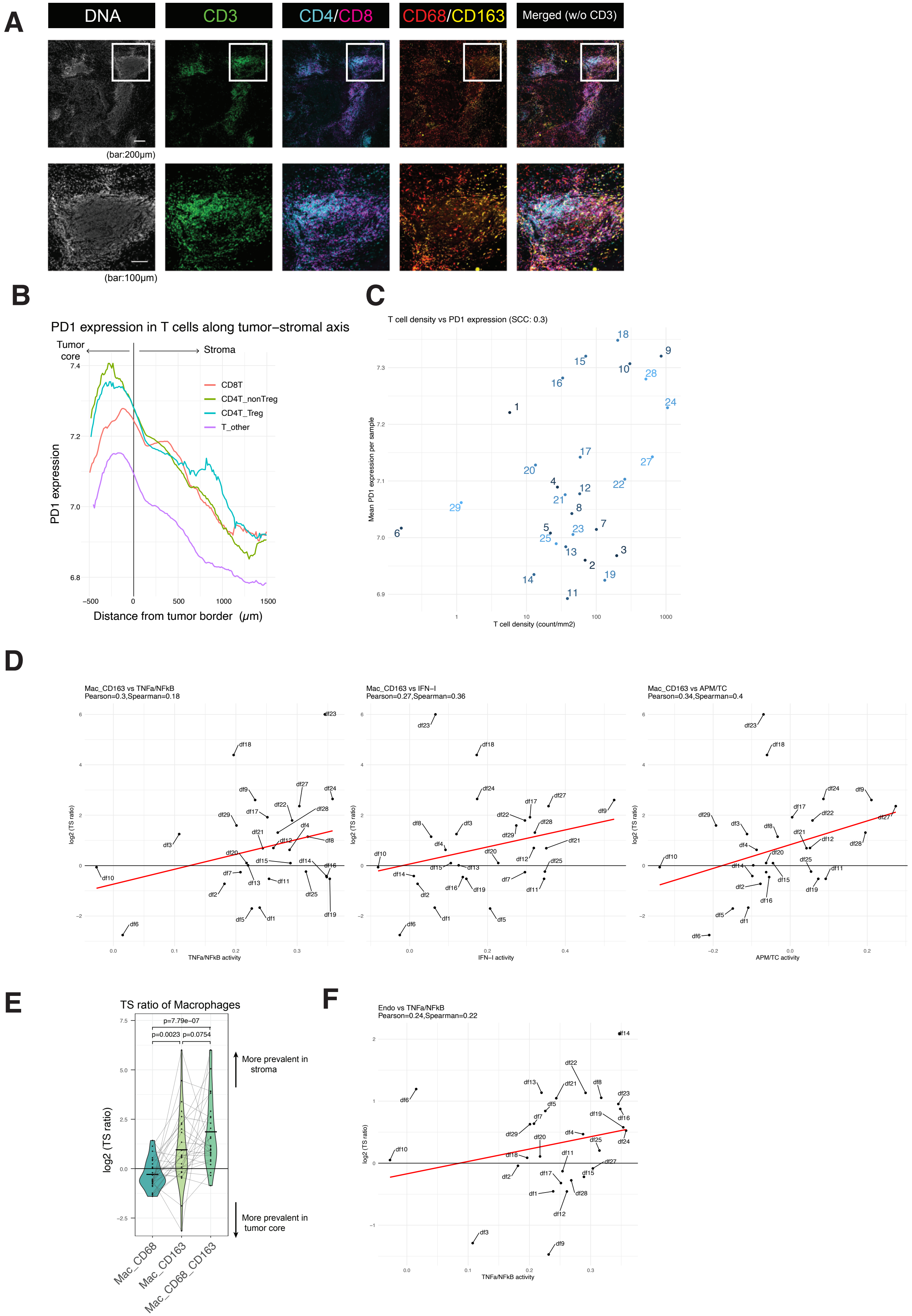

A
DNA
CD3
CD4/CD8
CD68/CD163
Merged (w/o CD3)
(bar:200μm)
(bar:100μm)
B
PD1 expression in T cells along tumor−stromal axis
Tumor core
Stroma
CD8T
CD4T_nonTreg
CD4T_Treg
T_other
PD1 expression
Distance from tumor border (μm)
C
T cell density vs PD1 expression (SCC: 0.3)
Mean PD1 expression per sample
T cell density (count/mm2)
D
Mac_CD163 vs TNFa/NFkB
Pearson=0.3,Spearman=0.18
TNFa/NFkB activity
Mac_CD163 vs IFN−I
Pearson=0.27,Spearman=0.36
IFN−I activity
Mac_CD163 vs APM/TC
Pearson=0.34,Spearman=0.4
APM/TC activity
log2 (TS ratio)
E
TS ratio of Macrophages
p=7.79e−07
p=0.0023
p=0.0754
More prevalent in stroma
More prevalent in tumor core
log2 (TS ratio)
Mac_CD68
Mac_CD163
Mac_CD68_CD163
F
Endo vs TNFa/NFkB
Pearson=0.24,Spearman=0.22
log2 (TS ratio)
TNFa/NFkB activity

### Figure. S10

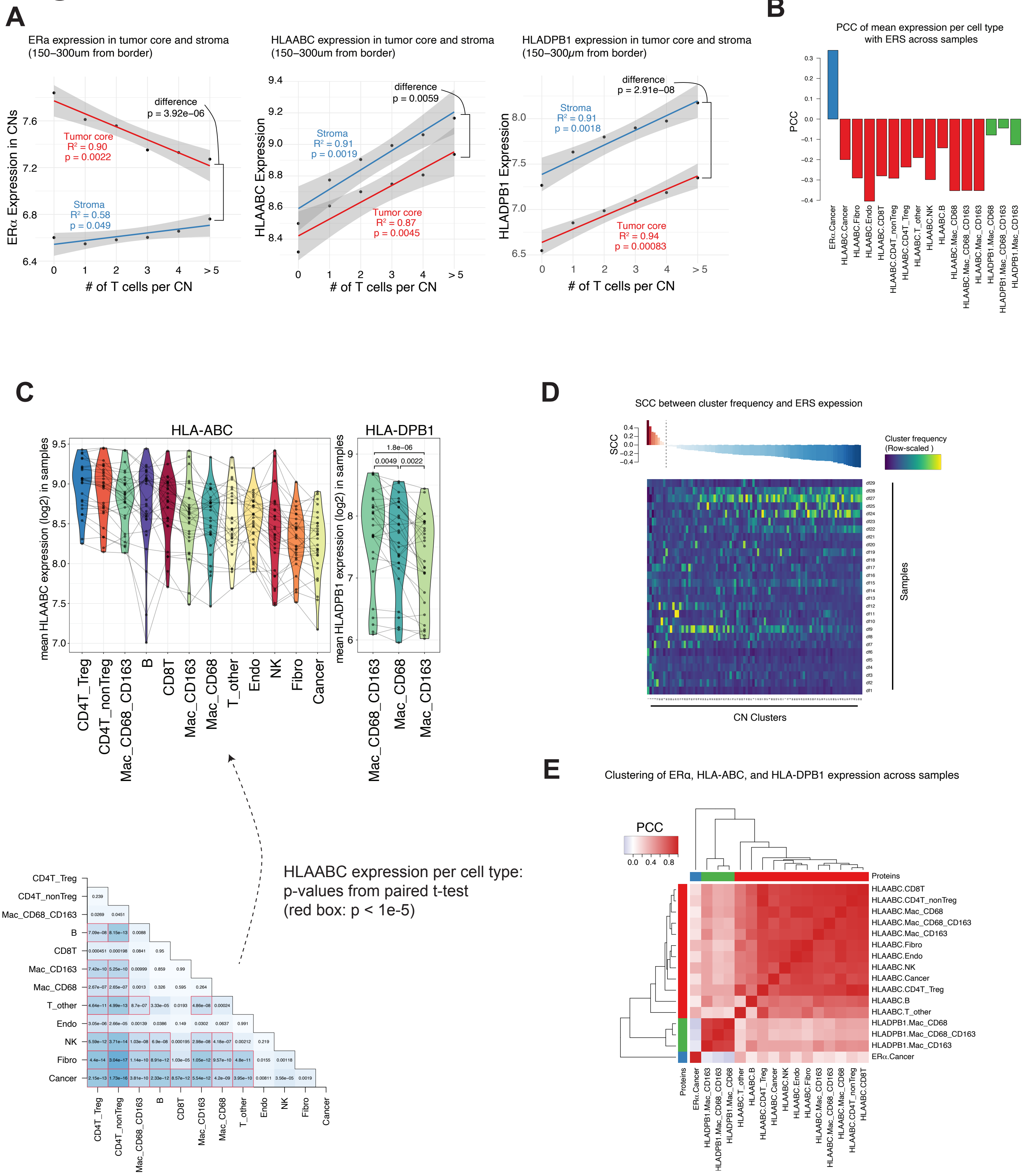
A
ERa expression in tumor core and stroma (150–300um from border)
ERα Expression in CNs
### of T cells per CN
Tumor core R² = 0.90 p = 0.0022
Stroma R² = 0.58 p = 0.049
difference p = 3.92e−06
HLAABC expression in tumor core and stroma (150–300um from border)
HLAABC Expression
Stroma R² = 0.91 p = 0.0019
Tumor core R² = 0.87 p = 0.0045
difference p = 0.0059
HLADPB1 expression in tumor core and stroma (150–300μm from border)
HLADPB1 Expression
Stroma R² = 0.91 p = 0.0018
Tumor core R² = 0.94 p = 0.00083
difference p = 2.91e−08
B
PCC of mean expression per cell type with ERS across samples
PCC
ERα.Cancer
HLAABC.Cancer
HLAABC.Fibro
HLAABC.Endo
HLAABC.CD8T
HLAABC.CD4T_nonTreg
HLAABC.CD4T_Treg
HLAABC.T_other
HLAABC.NK
HLAABC.B
HLAABC.Mac_CD68
HLAABC.Mac_CD68_CD163
HLAABC.Mac_CD163
HLADPB1.Mac_CD68
HLADPB1.Mac_CD68_CD163
HLADPB1.Mac_CD163
C
HLA-ABC
mean HLAABC expression (log2) in samples
CD4T_Treg
CD4T_nonTreg
Mac_CD68_CD163
B
CD8T
Mac_CD163
Mac_CD68
T_other
Endo
NK
Fibro
Cancer
HLA-DPB1
mean HLADPB1 expression (log2) in samples
1.8e−06
0.0049
0.0022
Mac_CD68_CD163
Mac_CD68
Mac_CD163
HLAABC expression per cell type: p-values from paired t-test (red box: p < 1e-5)
D
SCC between cluster frequency and ERS expression
SCC
Cluster frequency (Row-scaled )
Samples
CN Clusters
E
Clustering of ERα, HLA-ABC, and HLA-DPB1 expression across samples
PCC
0.0 0.4 0.8
Proteins
HLAABC.CD8T
HLAABC.CD4T_nonTreg
HLAABC.Mac_CD68
HLAABC.Mac_CD68_CD163
HLAABC.Mac_CD163
HLAABC.Fibro
HLAABC.Endo
HLAABC.NK
HLAABC.Cancer
HLAABC.CD4T_Treg
HLAABC.B
HLAABC.T_other
HLADPB1.Mac_CD68
HLADPB1.Mac_CD68_CD163
HLADPB1.Mac_CD163
ERα.Cancer

### Figure S11

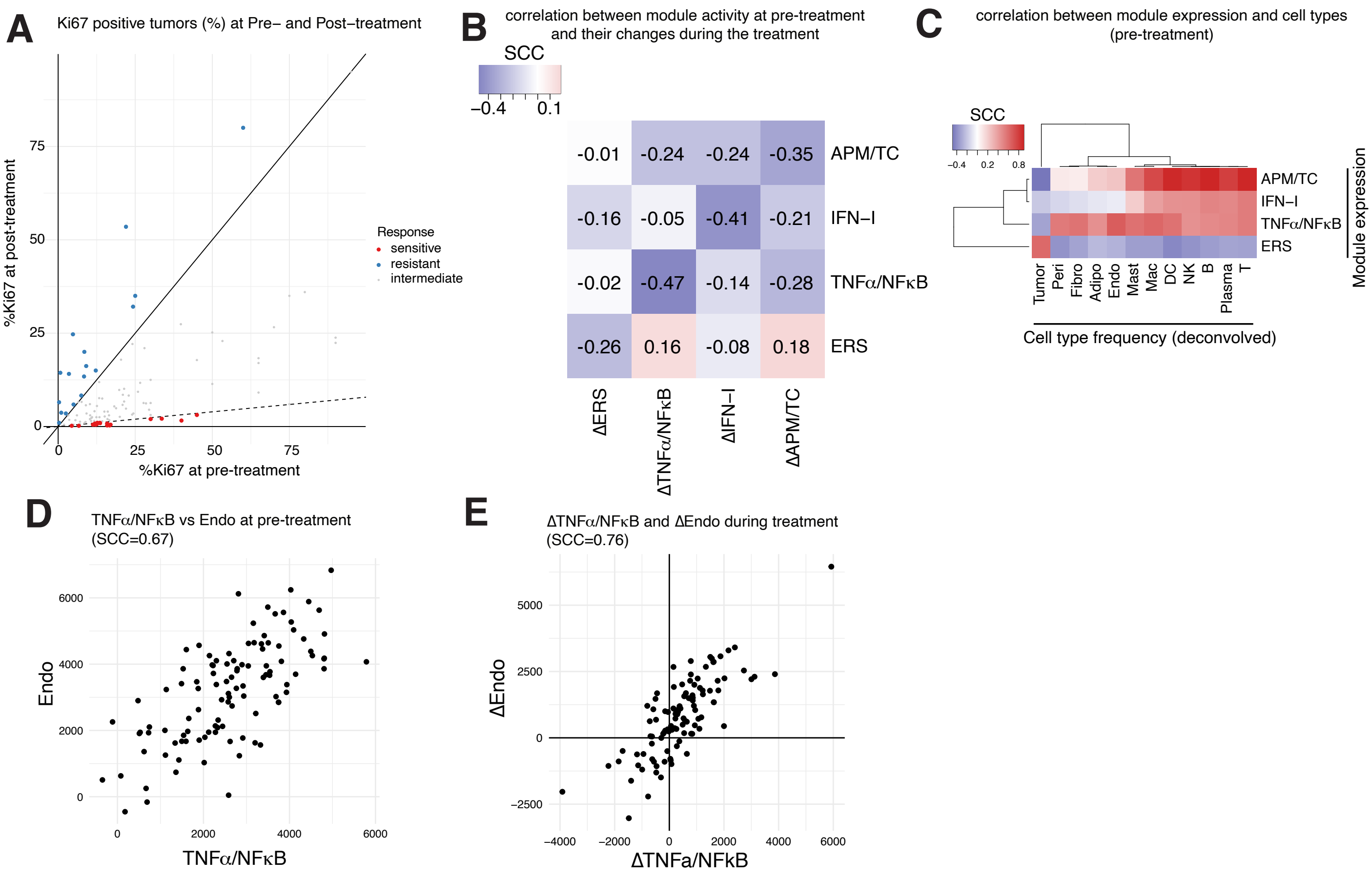
A
Ki67 positive tumors (%) at Pre– and Post–treatment
%Ki67 at post-treatment
%Ki67 at pre-treatment
Response
sensitive
resistant
intermediate
B
correlation between module activity at pre-treatment
and their changes during the treatment
SCC
–0.4
0.1
-0.01
-0.24
-0.24
-0.35
APM/TC
-0.16
-0.05
-0.41
-0.21
IFN–I
-0.02
-0.47
-0.14
-0.28
TNFα/NFκB
-0.26
0.16
-0.08
0.18
ERS
ΔERS
ΔTNFα/NFκB
ΔIFN–I
ΔAPM/TC
C
correlation between module expression and cell types
(pre-treatment)
SCC
APM/TC
IFN–I
TNFα/NFκB
ERS
Module expression
Tumor
Peri
Fibro
Adipo
Endo
Mast
Mac
DC
NK
B
Plasma
T
Cell type frequency (deconvolved)
D
TNFα/NFκB vs Endo at pre-treatment
(SCC=0.67)
Endo
TNFα/NFκB
E
ΔTNFα/NFκB and ΔEndo during treatment
(SCC=0.76)
ΔEndo
ΔTNFa/NFkB
